# Amplitude modulations of sensory responses, and deviations from Weber’s Law in pulsatile evidence accumulation

**DOI:** 10.1101/2020.06.24.167213

**Authors:** Sue Ann Koay, Stephan Y. Thiberge, Carlos D. Brody, David W. Tank

## Abstract

How do animals make behavioral decisions based on noisy sensory signals, which are moreover a tiny fraction of ongoing activity in the brain? Some theories suggest that sensory responses should be accumulated through time to reduce noise. Others suggest that feedback-based gain control of sensory responses allow small signals to be selectively amplified to drive behavior. We recorded from neuronal populations across posterior cortex as mice performed a decision-making task based on accumulating randomly timed pulses of visual evidence. Here we focus on a subset of neurons, with putative sensory responses that were time-locked to each pulse. These neurons exhibited a variety of amplitude (gain-like) modulations, notably by choice and accumulated evidence. These neural data inspired a hypothetical accumulation circuit with a multiplicative feedback-loop architecture, which parsimoniously explains deviations in perceptual discrimination from Weber-Fechner Law. Our neural observations thus led to a model that synthesizes both accumulation and feedback hypotheses.

## Introduction

As sensory information about the world is often noisy and/or ambiguous, an evidence accumulation process for increasing signal-to-noise ratio is thought to be fundamental to perceptual decision-making. Neural circuits that perform this are incompletely known, but canonically hypothesized to involve multiple stages starting from the detection of momentary sensory signals, which are then accumulated through time and later categorized into an appropriate behavioral action (Gold and Shadlen 2007; Brody and Hanks 2016; Caballero, Humphries, and Gurney 2018). In this picture, the sensory detection stage has a predominantly feedforward role, i.e. providing input to but not otherwise involved in accumulation and decision formation. However, another large body of literature has demonstrated that sensory processing in even the earliest sensory cortices can be modified by various external and internal contexts, including motor feedback, temporal statistics, learned associations, and attentional control (Roelfsema and de Lange 2016; Gilbert and Sigman 2007; Kimura 2012; Gavornik and Bear 2014; Glickfeld and Olsen 2017; Niell and Stryker 2010; Saleem et al. 2013; Shuler and Bear 2006; Fiser et al. 2016; Haefner, Berkes, and Fiser 2016; T. S. Lee and Mumford 2003; Zhang et al. 2014; Saleem et al. 2018; Makino and Komiyama 2015; Keller, Bonhoeffer, and Hü bener 2012; Poort et al. 2015; Li, Piëch, and Gilbert 2004; Stănişor et al. 2013; Petreanu et al. 2012; Romo et al. 2002; Luna et al. 2005; Nienborg, Cohen, and Cumming 2012; Yang et al. 2016; Britten et al. 1996; Froudarakis et al. 2019; Keller and Mrsic-Flogel 2018). For example, feedback-based gain control of sensory responses has been suggested as an important mechanism for enhancing behaviorally relevant signals, while suppressing irrelevant signals (Manita et al. 2015; Hillyard, Vogel, and Luck 1998; Harris and Thiele 2011; Azim and Seki 2019; Douglas and Martin 2007; Ahissar and Kleinfeld 2003).

The above two ideas—evidence accumulation and context-specific modulations—make two different but both compelling points about how sensory signals should be processed to support behavior. The two are not mutually incompatible. Here, we ask if a combination of both hypotheses could link neural to behavioral observations in a perceptual decision-making task. To observe how each sensory increment influences neural dynamics, we utilized a behavioral paradigm with precisely controlled timings of sensory inputs that should drive an evidence accumulation process (Brunton, Botvinick, and Brody 2013). Specifically, we recorded from posterior cortical areas during a navigational decision-making task (Pinto et al. 2018; BRAIN CoGS Collaboration, n.d.) where as mice ran down the central corridor of a virtual T-maze, pulses of visual evidence (“cues”) randomly appeared along both left and right sides of the corridor. To obtain rewards, mice should accumulate the numerosities of cues, then turn down the maze arm corresponding to the side with more cues. The well-separated and randomized timing of cues allowed us to clearly identify putative sensory responses that were time-locked to each pulse, while the seconds-long periods over which cues were delivered allowed us to observe the timecourse of neural responses throughout a gradually unfolding decision.

Across posterior cortices, the bulk of neural activity was sequentially active vs. time in the trial, in a manner that did not depend directly on the sensory cues, as we describe in detail in another article (Koay et al., n.d.). Even in the primary (V1) and secondary visual areas, only 5-10% of neurons had responses that were time-locked to sensory cues (“cue-locked cells”). Still, it is known that remarkably small signals on the order of a few cortical neurons can influence behavior (Doron and Brecht 2015; Buchan and Rowland 2018; Tanke, Borst, and Houweling 2018; Lerman et al. 2019; Carrillo-Reid et al. 2019; Marshel et al. 2019). Here we focused on the cue-locked cells, as candidates for momentary sensory inputs that may drive an accumulation and decision-making process. The responses of these cells to cues were well-described by a single impulse response function per neuron, but with amplitudes that varied across the many cue presentations. These cue-response amplitudes varied systematically across time in the trial as well as across trials depending on behavioral context, thus suggesting gain modulation effects potentially related to decision-making dynamics. Across posterior cortices and including as early as in V1, these variations in cue-response amplitudes contained information about multiple visual, motor, cognitive, and memory-related contextual variables. Notably, in all areas about 50% of cue-locked cells had response amplitudes that depended on the choice reported by the animal at the end of the trial, or depended on the value of the gradually accumulating evidence. Top-down feedback, potentially from non-sensory regions in which the choice is formed, has been proposed to explain choice-related effects in sensory responses (Britten et al. 1996; Romo et al. 2003; Nienborg and Cumming 2009; Yang et al. 2016; Bondy, Haefner, and Cumming 2018; Wimmer et al. 2015; Haefner, Berkes, and Fiser 2016). The dependence on accumulating evidence that we observed supports the hypothesis that this feedback may originate from an accumulator that itself eventually drives choice.

Inspired by our neural observations that the amplitudes of sensory responses depended on accumulated counts, we propose a multiplicative feedback-loop circuit model where accumulator feedback acts as a dynamic gain on the sensory input. Interestingly, this model can explain a psychophysical effect in the performance of pulse-based evidence accumulation tasks for both mice (Pinto et al. 2018) and rats (Scott et al. 2015). Specifically, the perceptual accuracy of rodents deviated from Weber-Fechner Law (Fechner 1966; Gallistel and Gelman 2000), which states that for a fixed ratio of right-to-left cue counts, the discriminability of the difference in counts should not depend on total (right plus left) counts. However, we found that the psychophysical data only followed this law at high total counts, and at low total counts instead exhibited a rise in performance vs. counts. We show that this trend can be explained as a competition between three classes of noise that have different dependencies on the number of pulses. These are: (1) noise associated with the comparison of two accumulators, which has a constant scale and therefore vanishing effect at high accumulator counts; (2) per-pulse sensory noise, the effect of which diminishes at high counts due to the central limit theorem; and (3) modulatory noise related to per-pulse gain fluctuations, which scales with the number of accumulated pulses and produces Weber-Fechner scaling at high counts. We found that modulatory noisy feedback from an accumulator can explain the experimentally-observed choice/count dependence of *sensory* unit responses. Moreover, the behavioral transition between low-counts and high-counts Weber-Fechner scaling regimes was best predicted by the feedback-loop model, compared to models without feedback.

In sum, our findings suggest that a behaviorally important feature of the accumulation process may be a gradual gain change in per-pulse sensory responses, as reflected in neural activity in as early as V1. Our multiplicative feedback-loop circuit model proposes a novel account of neural mechanisms that may underlie rodent psychophysics in pulsatile evidence accumulation tasks. A rich body of previous works have aimed to uncover the neural bases of Weber-Fechner scaling, some major ideas of which are that (a) the brain may represent the magnitude of sensory stimuli using a nonlinear (power law or logarithmic) scale (Fechner 1966; Dehaene and Changeux 1993; Nieder and Dehaene 2009); (b) perceptual accuracy may be limited by how evidence accumulation terminates upon reaching a bound (Pardo-Vazquez et al. 2019; Link 1992); and (c) noise at the memory level should dominate over noise at the sensory level (Gallistel and Gelman 2000). Intriguingly, our neural and behavioral observations differ from these previous hypotheses in ways that were illuminated by our use of pulsatile stimuli in the task design. (a) Since these pulses consisted of identical visual cues, the scalings of cue-locked responses that we observed were not purely of a sensory-magnitude origin, but instead reflected internal factors that evolved with the decision-making process. (b) Rodent behavior was influenced by all pulses in a trial, suggesting an effectively unbounded accumulation process (Brunton, Botvinick, and Brody 2013; Scott et al. 2015; Pinto et al. 2018) that nevertheless exhibits Weber-Fechner scaling at sufficiently high pulse counts. (c) At low total pulse counts, rodent performance deviated from Weber-Fechner’s law, which our models indicated was because noise at the memory level only dominated over sensory noise at sufficiently high pulse counts. Our neurophysiology-inspired model thus points to a circuit architecture with accumulator feedback gain onto sensory units as a potential cause of Weber-Fechner scaling that is distinct from (but may operate alongside) previous hypotheses (a-c).

## Results

We used cellular-resolution two-photon imaging to record from six posterior cortical regions of 11 mice trained in the Accumulating-Towers task (Fig. 1a-c). These mice were from transgenic lines that express the calcium-sensitive fluorescent indicator GCaMP6f in cortical excitatory neurons (Methods), and prior to behavioral training underwent surgical implantation of an optical cranial window centered over either the right or left parietal cortex. The mice then participated in previously detailed behavioral shaping (Pinto et al. 2018) and neural imaging procedures as summarized below.

**Figure 1.**
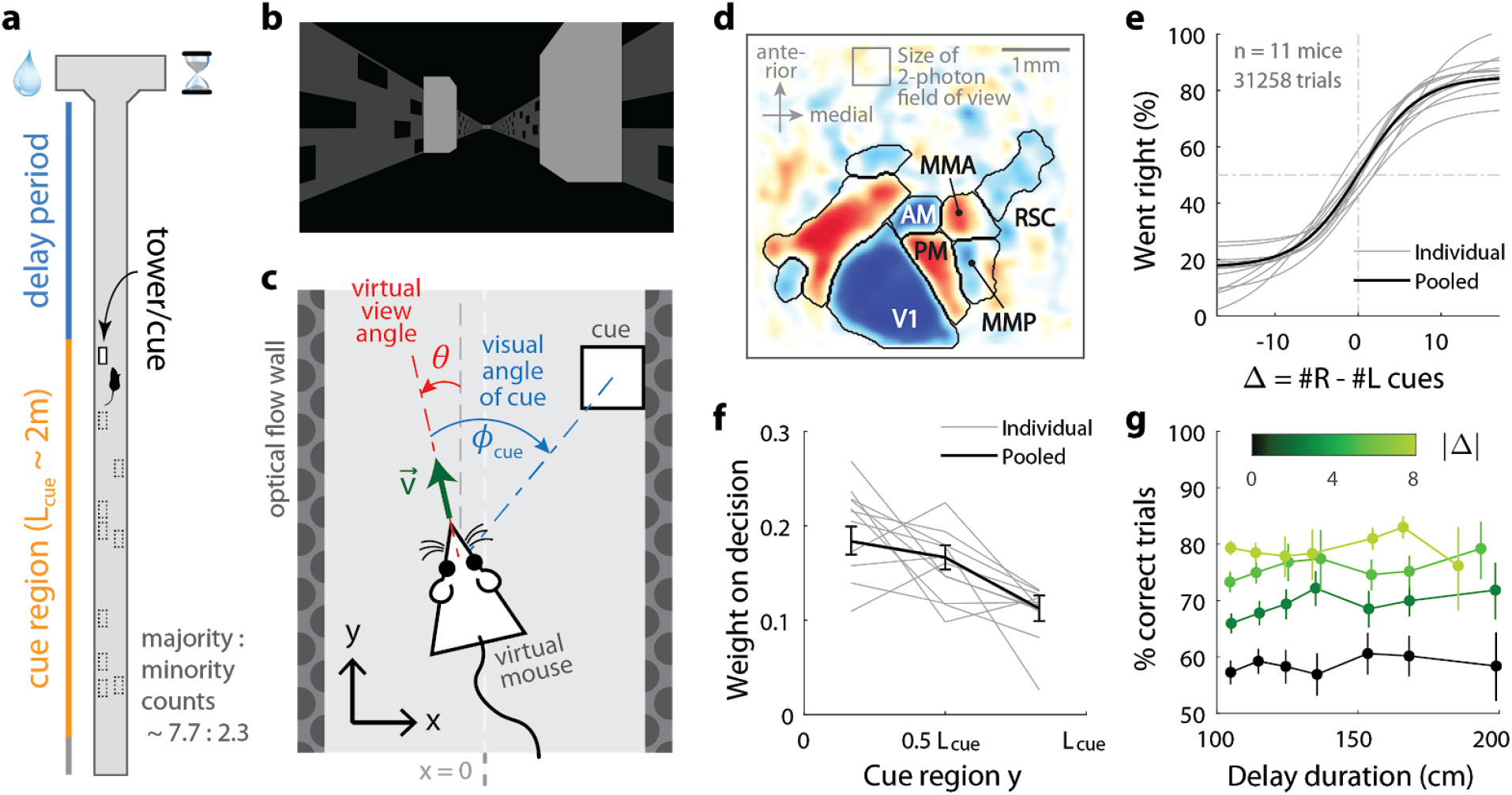
Two-photon calcium imaging of posterior cortical areas during a navigation-based evidence accumulation task. **(a)** Layout of the virtual T-maze in an example left-rewarded trial. **(b)** Example snapshot of the cue region corridor from a mouse’s point of view when facing straight down the maze. Two cues on the right and left sides can be seen, closer and further from the mouse in that order. **(c)** Illustration of the virtual viewing angle θ. The visual angle φ_*cue*_ of a given cue is measured relative to θ and to the center of the cue. The *y* spatial coordinate points straight down the stem of the maze, and the *x* coordinate is transverse. 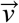 is the velocity of the mouse in the virtual world. **(d)** Average visual field sign map (*n* = 5 mice) and visual area boundaries, with all recorded areas labeled. The visual field sign is −1 (dark blue) where the cortical layout is a mirror image and +1 (dark red) where it follows a non-inverted layout of the physical world. **(e)** Sigmoid curve fits to behavioral data for how frequently mice turned right for a given difference in total right vs. total left cue counts at the end of the trial, Δ ≡ #*R*−#*L*. **(f)** Logistic regression weights for predicting the mice’s choice given spatially-binned evidence {Δ_*i*_} where *i* ∈ {1, 2, 3} indexes three equally-sized spatial bins of the cue region. Error bars: 95% C.I. across bootstrap experiments. **(g)** Performance vs. effective duration of the delay period, which is the distance from the last cue to the end of the T-maze stem. Data were pooled over trials with all total cue counts #*R* + #*L* for statistical power, but analyses that account for #*R* + #*L* differences yield the same conclusion (Pinto et al. 2018). Error bars: 95% C.I. across trials within various bins of Δ ≡ #*R* − #*L* (data from all sessions).

Mice were trained in a head-fixed virtual reality system (Dombeck et al. 2010) to navigate in a T-maze. As they ran down the stem of the maze, a series of transient (200ms), randomly located tower-shaped cues (Fig. 1b) appeared along the right and left walls of the cue region corridor (length *L*_*cue*_ ≈ 200 cm, average running speed in cue region ≈ 60*cm*/*s*; see Methods), followed by a delay region where no cues appeared. The locations of cues were drawn randomly per trial, with Poisson-distributed mean counts of 7.7 on the majority and 2.3 on the minority side, and mice were rewarded for turning down the arm corresponding to the side with more cues. As mice control the virtual viewing angle θ, cues could appear at a variety of visual angles φ_*cue*_ (Fig. 1c). We accounted for this in all relevant data analyses, as well as conducted control experiments in which θ was restricted to be exactly zero from the beginning of the trial up to midway in the delay period (referred to as θ-controlled experiments; see Methods). In agreement with previous work (Pinto et al. 2018), all mice in this study exhibited characteristic psychometric curves and utilized multiple pieces of evidence to make decisions, with a small primacy effect (Fig. 1e-f). For fixed total numbers of cues on the right (#R) and left (#L) sides, there was no degradation of performance with increasing effective length of the delay period (Fig. 1g). This is compatible with a negligible effect of memory leakage with time, as observed also in rats performing evidence-accumulation tasks (Brunton, Botvinick, and Brody 2013; Scott et al. 2015).

For each mouse, we first identified the locations of the visual areas (Fig. 1d; Methods) using one-photon widefield imaging and a retinotopic visual stimulation protocol (Zhuang et al. 2017). Then, while the mice performed the task, we used two-photon imaging to record from 500μ*m* × 500μ*m* fields of view in either layers 2/3 or 5 from one of six areas (Supplementary Table 1, Supplementary Table 2): the primary visual cortex (V1), secondary visual areas (V2 including AM, PM, MMA, MMP (Zhuang et al. 2017)), and retrosplenial cortex (RSC). After correction for rigid brain motion, regions of interest representing putative single neurons were extracted using a semi-customized (Methods) demixing and deconvolution procedure (Pnevmatikakis et al. 2016). The fluorescence-to-baseline ratio Δ*F* /*F* was used as an estimator of neural activity, and only cells with ≥ 0.1 transients per trial were selected for analysis. In total, we analyzed 10,481 cells from 145 imaging sessions.

### Pulses of evidence evoke transient, time-locked responses in all recorded areas

We found neurons in all areas/layers that had activities clearly time-locked to the pulsatile cues (examples in Fig. 2a-b). In trials with sparse occurrences of preferred-side cues, the activities of these cells tended to return to baseline following a fairly stereotyped impulse response. Individually, they thus represented only momentary information about the visual cues, although as a population they can form a more persistent stimulus memory (Goldman 2009; Scott et al. 2017; Miri et al. 2011). Interestingly, the amplitudes of these cells’ responses seemed to vary in a structured way, both across time in a trial, as well as across trials where the mouse eventually makes the choice to turn right vs. left (columns of Fig. 2a-b). We therefore wished to quantify whether or not these putatively sensory amplitude changes also encoded other task-related information.

**Figure 2.**
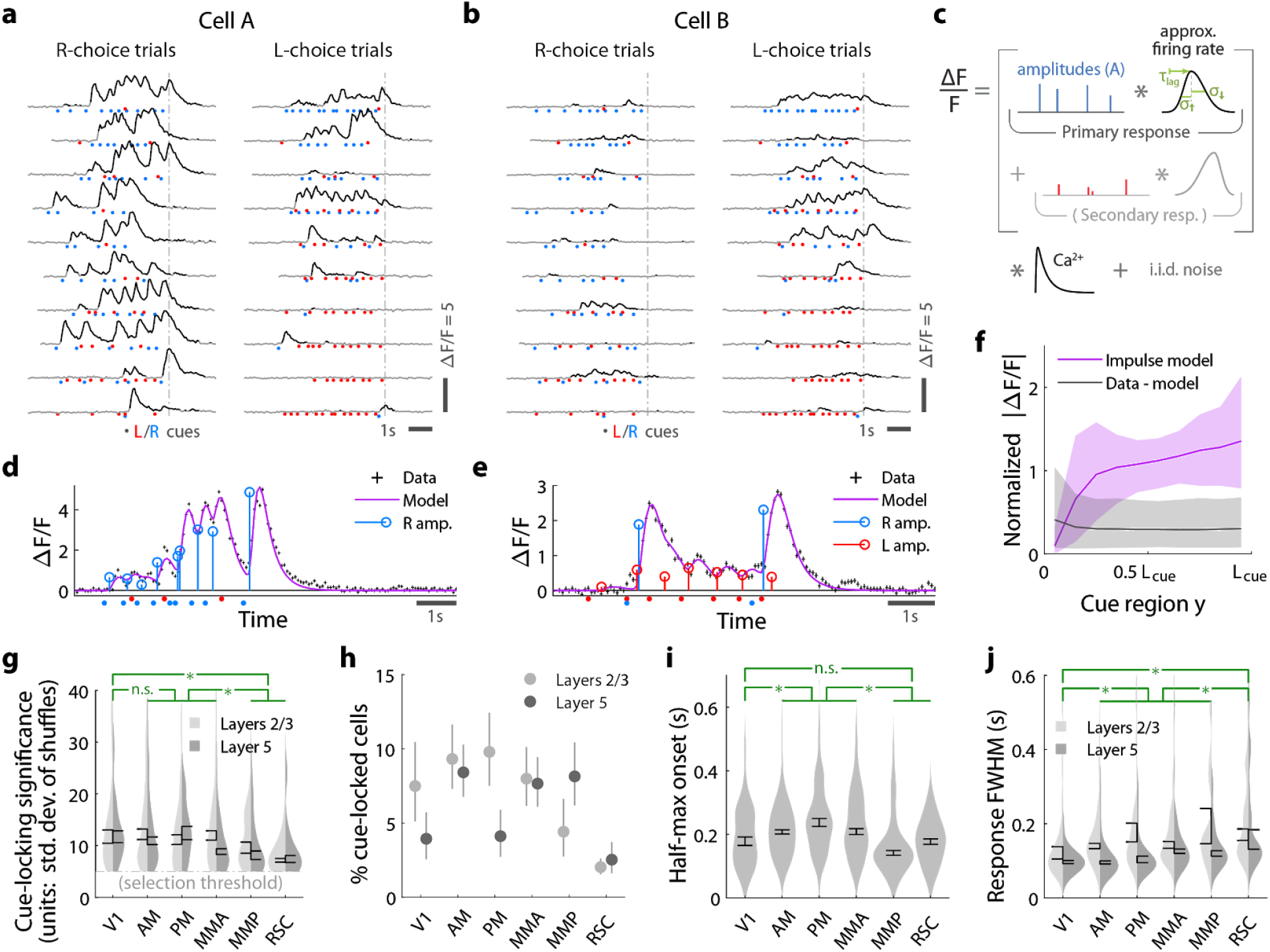
Pulses of evidence evoke transient, time-locked responses that are well described by an impulse response model. **(a)** Trial-by-trial activity (rows) vs. time of an example right-cue-locked cell recorded in area AM, aligned in time to the end of the cue period (dashed line). Onset times of left (right) cues in each trial are shown as red (blue) dots. **(b)** Same as (a), but for an *atypical* right-cue-locked cell (in area AM) that has some left-cue-locked responses. **(c)** Depiction of the impulse response model for the activity level Δ*F* /*F* of a neuron vs. time (x-axis). Star indicates the convolution operator. **(d)** Prediction of the impulse response model for the cell in (a) in one example trial. This cell had no significant secondary (left-cue) responses. **(e)** Same as (d) but for the cell in (b). The model prediction is the sum of primary (right-cue) and secondary (left-cue) responses. **(f)** Trial-average impulse response model prediction (purple) vs. the residual of the fit (Δ*F* /*F* data minus model prediction, black), in 10 equally-sized spatial bins of the cue region. For a given cell, the average model prediction (or average residual) is computed in each spatial bin, then the absolute value of this quantity is averaged across trials, separately per spatial bin. Line: Mean across cells. Band: 95% C.I. across cells. For comparability across cells, Δ*F* /*F* was expressed in units such that the mean model prediction of each cell is 1. The model prediction rises gradually from baseline at the beginning of the cue period due to nonzero lags in response onsets. **(g)** Distribution (kernel density estimate) of cue-locking significance for cells in various areas/layers. Significance is defined per cell, as the number of standard deviations beyond the median AIC_C_ score of models constructed using shuffled data (Methods). Error bars: S.E.M. of cells. Stars: significant differences in means (Wilcoxon rank-sum test). **(h)** Percent of significantly cue-locked cells in various areas/layers. Chance: 3 × 10^−5^%. Error bars: 95% binomial C.I. across sessions. **(i)** Distribution (kernel density estimate) of the half-maximum onset time of the primary response, for cells in various areas. Data were pooled across layers (inter-layer differences not significant). Error bars: S.E.M. across cells. Stars: significant differences in means (Wilcoxon rank-sum test). **(j)** As in (i) but for the full-width-at-half-max. Statistical tests use data pooled across layers. Means were significantly different across layers for all areas except V1 and RSC (Wilcoxon rank-sum test).

For a given cell, we estimated the amplitude of its response to each cue *i* by modeling the cell’s activity as a time series of non-negative amplitudes *A*_*i*_ convolved with an impulse response function (Fig. 2c). The latter was defined by lag, rise-time and fall-time parameters that were fit to each cell, but were the same for all cue-responses of that cell (deconvolving calcium dynamics; see Methods). For a subset of neurons, this impulse response model resulted in excellent fits when the model included only primary responses to either right- or left-side cues (e.g. Fig. 2d). In much rarer instances, adding a secondary response to the opposite-side cues resulted in a significantly better fit (e.g. Fig. 2e; discounting for number of parameters by using AIC_C_ (Hurvich and Tsai 1989) as a measure of goodness of fit). We defined cells to be cue-locked if the primary-response model yielded a much better fit to the data than a permutation test (data with cue timings shuffled within the cue region, see Methods). For these cells, the trial-averaged activity predicted by the impulse response model (Fig. 2f, “signal” in magenta) was substantially larger than the magnitude of residuals of the fits (Fig. 2f, “data – model prediction” in black). For example, if cells had systematic rises or falls in baseline activity levels vs. time/place that could not be explained as transient responses to cues, then the residual would grow/diminish vs. *y* location in the cue region. Supplementary Fig. 1a shows that systematic trends (i.e. slopes) for the residual vs. *y* was small for most cells (68% C.I. of slopes across cells were within [− 0.063, 0.042], where a slope of ± 1 corresponds to a change in residuals from the start to the end of the cue region being equal to the average signal predicted by the impulse response model). There were thus no large, unaccounted-for components in the activity of these identified cue-locked cells, in particular no components with long timescales.

Significantly cue-locked cells comprised a small fraction of the overall neural activity, but were nevertheless present in all areas/layers and exhibited some progression of response properties across posterior cortical areas in a roughly lateral-to-medial order (V1, V2, RSC). Cells with the most precisely time-locked responses to cues were found in the visual areas as opposed to RSC (high-significance tail of distributions in Fig. 2g; low significance means that the model fit comparably well to data where cue timings were shuffled within the cue region). Reflecting this, about 5-10% of cells in visual areas were significantly cue-locked, compared to ~ 2% in RSC (Fig. 2h). Of these significant cells, only ~ 5% had secondary responses that were moreover much less significantly time-locked (Supplementary Fig. 1b); most cells responded to only contralateral cues (Supplementary Fig. 1c). The onset of the half-maximum response was ~ 200*ms* after each pulse (Fig. 2i), and the response full-width-at-half-max (FWHM) was ~ 100*ms* but increased from V1 to secondary visual areas to RSC (Fig. 2j). The impulse response model thus identified cells that follow what one might expect of purely visual-sensory responses on a cue-by-cue basis, but up to amplitude changes that we next discuss.

### Cue-locked response amplitudes contain information about visual, motor, cognitive, and memory-related contextual task variables

Studies of perceptual decision-making have shown that the animal’s upcoming choice affects the activity of stimulus-selective neurons in a variety of areas (Britten et al. 1996; Nienborg and Cumming 2009). We analogously looked for such effects (and more) while accounting for the highly dynamical nature of our task. As neurons responded predominantly to only one laterality of cues, all our subsequent analyses focus on the primary-response amplitudes of cue-locked cells. Importantly, the impulse response model deconvolves responses to individual cues, so the response amplitude *A*_*i*_ can be conceptualized as a multiplicative gain factor that the cell’s response was subject to at the instant at which the *i*^*th*^ cue appeared.

We used a neural-population decoding analysis to quantify how much information the cue-locked response amplitudes contained about various contextual variables. First, for the *i^th^* cue in the trial, we defined the neural state as the vector of amplitudes *A*_*i*_ of cells that responded to contralateral cues only. Then using the neural states corresponding to cues that occurred in the first third of the cue period, we trained a support vector machine (SVM) to linearly decode a given task variable from these neural states (cross-validated and corrected for multiple comparisons; see Methods). This procedure was repeated for the other two spatial bins (second third and final third) of the cue period, to observe changes in neural information that may reflect place-/time-related changes in task conditions (illustrated in Fig. 3a). Fig. 3b shows that across posterior cortex, four task variables were accurately decodable from the cue-response amplitudes: the view angle θ, running speed, the running tally of evidence (Δ ≡ #*R* − #*L*), and the eventual choice to turn right or left. The reward outcome from the previous trial could also be decoded, albeit less accurately, while in contrast decoding of the past-trial choice was near chance levels.

**Figure 3.**
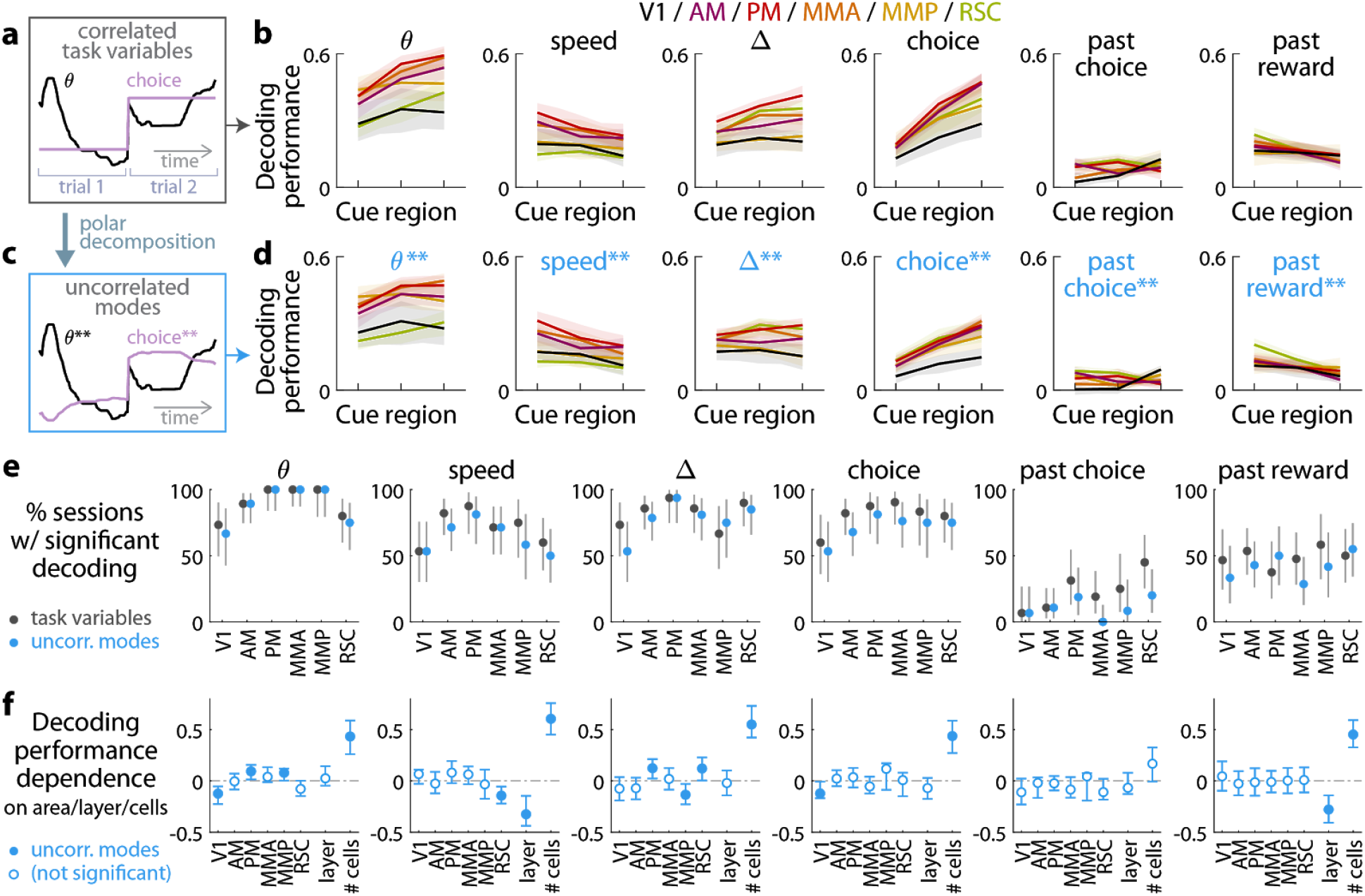
Multiple visual, motor, cognitive, and memory-related variables can be decoded from the amplitudes of cue-locked cell responses. **(a)** Example time-traces of two statistically correlated task variables, the view angle θ (black) and the eventual navigational choice (magenta). **(b)** Cross-validated performance for decoding six task variables (individual plots) from the amplitudes of cue-locked neuronal responses, separately evaluated using responses to cues in three spatial bins of the cue region (Methods). The performance measure is Pearson’s correlation between the actual task variable value and the prediction using cue-locked cell amplitudes. Lines: mean performance across recording sessions for various areas (colors). Bands: S.E.M. across sessions, for each area. **(c)** Example time-traces of the two uncorrelated modes obtained from a polar decomposition of the correlated task variables in (a). This decomposition (Methods) solves for these uncorrelated modes such that they were linear combinations of the original time-traces that were closest, in the least-squares sense, to the original traces, while constrained to be themselves uncorrelsome example cells witha ted with each other. Correlation coefficients between individual uncorrelated modes and their corresponding original variables were > 0.85 for all modes (Supplementary Fig. 2). **(d)** As in (a), but for decoding the uncorrelated task-variable modes illustrated in (c). **(e)** Proportion of imaging sessions that had significant decoding performance for the six task variables in (b) (dark gray points) and uncorrelated modes in (d) (blue points), compared to shuffled data and corrected for multiple comparisons. Data were restricted to 117 sessions with at least one cue-locked cell. Error bars: 95% binomial C.I. across sessions. **(f)** Linear regression (Support Vector Machine) weights for how much the decoding performance for uncorrelated task-variable modes in (d) depended on cortical area/layer and number of recorded cue-locked cells. Weights that are not statistically different from zero are indicated with open circles. The negative weight for layer dependence of past-reward decoding means that layer 5 had significantly lower decoding performance than layers 2/3. Error bars: 95% C.I. computed via bootstrapping sessions.

As the six variables were statistically correlated by nature of the task (e.g. the mouse controls θ to execute the navigational choice, Fig. 3a), indirect neural information about one variable could be exploited to increase the performance of decoding another correlated variable (Krumin et al. 2018; Koay et al., n.d.). To account for this, we repeated the decoding analyses for a modified set of variables that had statistical correlations removed. As explained in the Methods and illustrated in Fig. 3c, we solved for uncorrelated modes being linear combinations of the original time-traces that were closest, in the least-squares sense, to the original traces, while constrained to be themselves uncorrelated with each other. As inter-variable correlations were low throughout the cue region, these uncorrelated modes were very similar to the original task variables. Each uncorrelated mode was identified with its closest original variable and labeled as such, and correlation coefficients between individual uncorrelated modes and their corresponding original variables were > 0.85 for all modes (Supplementary Fig. 2). Performances for decoding the uncorrelated modes were a little lower than the original task variables (Fig. 3d), as expected since contributions from indirect neural information could no longer be present. Nevertheless, the modes that resembled view angle, speed, evidence, choice, and past-trial reward could all be consistently decoded across imaging sessions for all examined areas (Fig. 3e). There was also comparably high performance of decoding evidence and choice in the θ–controlled experiments (Supplementary Fig. 3), which explicitly shows that neural information about these variables do not originate solely from changes in visual perspective. In a comparable task where choice was highly correlated with view angle (θ) and *y* spatial location in the maze, it has previously been reported that θ and *y* explains most of neural responses in parietal posterior cortex, with small gains from including choice as a third factor (Krumin et al. 2018). Interestingly however, our findings indicate that in a task where choice was *distinguishable* from other behavioral factors (here, at least within the cue region), there was significant neural information in all surveyed posterior cortical areas about this internally generated variable, choice.

As a population, the amplitudes of cue-locked cells thus reflected a rich set of present- and past-trial contextual information, with some anatomical differences. Likely due to the small numbers of recorded cue-locked cells per session (~0-10), the decoding performance for all variables depended most strongly on the number of cells (Fig. 3f). Fig. 3f also shows that V1 had a small but significantly lower view angle and choice decoding performance than other posterior parietal regions, whereas RSC had significantly lower speed decoding and higher evidence decoding performance than other regions. Layer 5 was also distinguishable from layer 2/3 data in having reduced performance for decoding speed and past-trial reward.

### Decision-related changes in cue-locked response amplitudes are compatible with a feedback origin

Interestingly, the response amplitudes of some individual cue-locked cells appeared to systematically depend on time (e.g. Fig. 2a-b), as did the population-level decoding performance for variables such as choice (Fig. 3b,d). To understand if these neural dynamics may reflect a gradually unfolding decision-making process, we turned to modeling how amplitudes of cue-locked cell responses may depend on choice and place/time, while accounting for other time-varying behavioral factors.

As a null hypothesis based on previous literature, we hypothesized that cue-response amplitudes can depend on a receptive field specified by the visual angle of the cue (φ_*cue*_, Fig. 1c), as well as running speed (Niell and Stryker 2010; Saleem et al. 2013). Given limited data statistics, we compared this null hypothesis to three other conceptually distinct models (Methods), each of which aims to parsimoniously explain cue-response amplitudes using small sets of behavioral factors. These models predict the observed cue-response amplitudes to be random samples from a Gamma distribution, where the mean of the Gamma distribution is a function of various behavioral factors at the time at which a given cue appeared. The mean functions for all models have the form ρ(φ_*cue*_) *f* (*v*) *g*(· · ·), where ρ(φ_*cue*_) is an angular receptive field function, *f* (*v*) is a running speed (*v*) dependence function, and *g*(· · ·) is specific to each of the three models, as follows. First, the “SSA” model parameterizes stimulus-specific adaptation (Ulanovsky, Las, and Nelken 2003; Sobotka and Ringo 1994) or enhancement (Vinken, Vogels, and Op de Beeck 2017; Kaneko, Fu, and Stryker 2017) with exponential time-recovery in between cues. Second, the “choice” model allows for a flexible change in amplitudes vs. place/time in the cue region, with a potentially different trend for right- vs. left-choice trials. Third, the “cue-counts” model allows the amplitudes to depend on the running tally of #*R*, #*L*, or Δ = #*R* − #*L*. This selection of models allows us to ask if cue-locked responses are sufficiently explained by previously known effects, or if after accounting for such there are still effects related to the accumulation process, such as choice or cue-count dependence.

We constructed the amplitude model prediction as the AIC_C_-likelihood-weighted average of the above models, which accounts for when two or more are comparably good (Volinsky et al. 1999). As illustrative examples, Fig. 4a shows how the amplitudes of two simultaneously recorded cue-locked cells in area AM depended on behavioral factors and compared to model predictions. There are clear differences in predictions for right- vs. left-choice trials that can also be seen in the raw amplitude data (restricted to a range of φ_*cue*_ such that angular receptive field effects are small, 2nd and 3rd columns of Fig. 4a). Although both cells responded preferentially to right-side cues, they had oppositely signed choice modulation effects, defined as the difference between amplitude model predictions on contralateral- vs. ipsilateral-choice trials (Methods). Fig. 4b shows two more example choice-modulated cells that had near-constant angular receptive fields. We note that except for the parameters of SSA, all findings in this section were qualitatively similar in θ-controlled experiments where there can be no angular receptive field effects (Supplementary Fig. 4).

**Figure 4.**
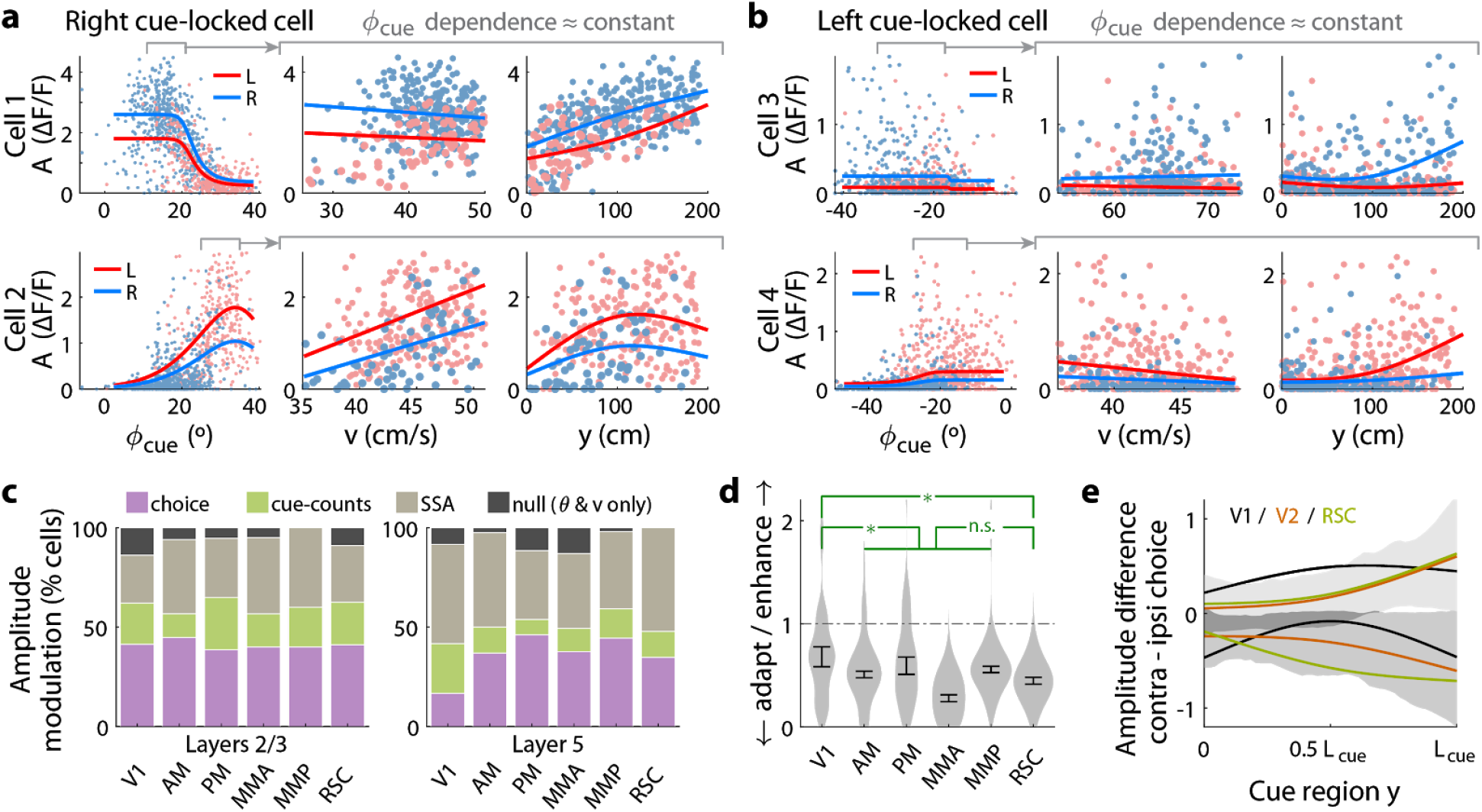
Cue-locked response amplitudes depend on view angle, speed, and cue frequency, but a large fraction exhibit choice-related modulations that increase during the course of the trial. **(a)** Response amplitudes of two example right-cue-locked cells (one cell per row) vs. (columns) the visual angle at which the cue appeared (φ_*cue*_), running speed (*v*), and *y* location of the cue in the cue region. Points: amplitude data in blue (red) according to the upcoming right (left) choice. Lines: AIC_C_-weighted model mean functions for right- vs. left-choice trials (lines); the model predicts the data to be random samples from a Gamma distribution with this behavior-dependent mean function. The data in the right two columns were restricted to a subset where angular receptive field effects are small, corresponding to the indicated area in the leftmost plots. **(b)** Same as (a) but for two (left-cue-locked) cells with broader angular receptive fields. **(c)** Percentages of cells that significantly favor various amplitude modulation models (likelihood ratio < 0.05, defaulting to null model if none are significant), in the indicated cortical areas and layers. For layer 2/3 data, PM vs. MMA had significantly different fractions of choice-model preferring cells (*p* = 0.03, two-tailed Wilcoxon rank-sum test). For layer 5 data, V1 has a significantly lower choice-model preferring fraction than the other areas (*p* = 0.033). **(d)** Distribution (kernel density estimate) of adaptation/enhancement factors for cells that favor the SSA model. A factor of 1 corresponds to no adaptation, while for other values the subsequent response is scaled by this amount with exponential recovery towards 1. Error bars: S.E.M. Stars: significant differences in means (Wilcoxon rank-sum test). **(e)** Choice modulation strength vs. location in the cue region, for contralateral-cue-locked cells only (see ipsilateral-cue-locked cells in Supplementary Fig. 5g). Lines: mean across cells in a given brain area (pooled across layers). Bands do not indicate significance but instead indicate distribution: they represent the std. dev. of data pooled across all areas/layers. Both mean and std. dev. are shown separately for cells with positive vs. negative modulations.

To summarize the prevalence and composition of amplitude-modulation effects, we selected the best model per cell using AIC_C_, defaulting in ambiguous cases (relative likelihood < 0.05) to the null hypothesis. Supplementary Fig. 5a shows that there were large fractions of cells with very high AIC_C_ likelihoods for all three alternative models compared to the null hypothesis. Cells that favored the cue-counts model could also be clearly distinguished from those that favored the SSA model (Supplementary Fig. 5b); in fact, exclusion of cells that exhibited SSA had little effect on how well evidence and other variables could be decoded from the neural population (Supplementary Fig. 6). In all areas and layers, > 86% of cue-locked cells exhibited some form of amplitude modulations beyond angular receptive field and running speed effects (Fig. 4c). Overall, 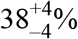 of cells were best explained by SSA while 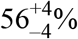 favored either choice or cue-counts models. The only notable inter-area differences is for layer 5 data, which had a smaller proportion of choice-model preferring cells in V1 compared to other areas (*p* = 0.03, Wilcoxon rank-sum test). Most cells thus exhibited some form of amplitude modulations beyond visuomotor effects, with little difference in composition across areas and layers.

Although SSA, choice, and cue-counts dependencies all predict changes in cue-response amplitudes vs. time in the trial, there were qualitative differences that distinguished SSA from choice and cue-count modulations, as we next discuss. Cells in the two largest categories, SSA and choice, had qualitatively different population statistics for how their cue-response amplitudes depended on place/time in the trial. Most cells 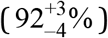 that favored the SSA model corresponded to a phenotype with decreased responses to subsequent cues. Adaptation effects were weakest in V1 and stronger other areas (Fig. 4d, but see Supplementary Fig. 4f-g for θ-controlled experiments), although the ~ 0.8*s* recovery timescale had no significant inter-area differences (Supplementary Fig. 5d). In contrast, cue-locked cells with both choice laterality preferences were intermixed in all areas and layers (Supplementary Fig. 5e). Also unlike the *decrease* in response amplitudes vs. time for cells that favored the SSA model, both subpopulations of positively and negatively choice-modulated cells exhibited gradually *increasing* effect sizes vs. place/time in the trial (Fig. 4e for contralateral cue-locked cells, Supplementary Fig. 5g for ipsilateral cue-locked cells). Cells that favored the cue-counts modulated category also had qualitatively different population statistics compared to cells that exhibited SSA. Comparable proportions of cue-counts modulated cells were best explained by dependence on counts on either the contralateral side, ipsilateral side, or the difference of the two sides (Supplementary Fig. 5f). For (say) right-cue-locked cells, #*L* or Δ dependencies are not directly explainable by SSA because of the modulation by left-side cues that the cells do not otherwise respond to. The remaining time-independent #*R* modulation also cannot be explained by SSA, unless SSA has an infinitely long timescale. Such infinite-timescale SSA would require some additional prescription for “resetting” the adaptation factor e.g. at the start of each trial, because otherwise amplitudes would continue to decrease/increase throughout the ~1 hour long session (which we do not observe).

Although relationships between sensory responses and choice can arise in a purely feedforward circuit structure, because sensory neurons play a causal role in producing the behavioral choice (Shadlen et al. 1996), others have noted that this should result in similar timecourses of neural and behavioral fluctuations (Nienborg and Cumming 2009). Instead, we observed contrasting timecourses: as each trial evolved, there was a slow *increase* in time in choice modulations of cue-locked responses (Fig. 4e; Supplementary Fig. 5g), which was opposite to the behaviorally-assessed *decrease* in time in how sensory evidence fluctuations influenced the mice’s choice (Fig. 1f). Additionally, a feedforward structure predicts that positive fluctuations in right- (left)-preferring cue-locked neurons should produce rightwards (leftwards) fluctuations in choice. Instead, we observed that about half of the cue-locked cells were modulated by choice in a manner opposite to their cue-side preference (Supplementary Fig. 5e). Both of these observations argue against a purely feedforward structure, and thus support the existence of feedback influences on sensory responses (Wimmer et al. 2015; Nienborg and Cumming 2009; Haefner, Berkes, and Fiser 2016).

### Cue-locked amplitude modulations motivate a multiplicative feedback-loop circuit model

We hypothesized that the cue-locked sensory responses in visual areas provide momentary cue information that drives an accumulation process, which ultimately drives choice. If this is true, modulations of cue-response amplitudes may predict specific perceptual biases in decision making. As previously argued, the observed timecourse of choice modulations point to a feedback effect from a decision-making process (Fig. 4e). The accumulation hypothesis would naturally propose that these effects originate from an evidence accumulator. Thus inspired by our physiological recordings, we propose that feedback projections from a pulse accumulator induce choice- and pulse-count-related modulations of cue-locked sensory responses. We used a simplified dynamical systems model to ask what behavioral consequences would be predicted by such feedback. Previous psychophysical analyses have suggested that there are two near-independent accumulators, one for the right stimulus stream, and another for the left, which are then compared to form the behavioral decision (Scott et al. 2015). Our model consequently has separate accumulators for the right and left stimulus streams. Below we describe the model for a right-side-specific accumulator given by a single scalar variable *a*_*r*_(*t*); identical results hold for a left-accumulator *a*_*l*_(*t*). We first discuss the solution of this mathematical model, address the impact of noise or stochasticity on the model, and show that it is qualitatively compatible with trends observed in the neural data. In the next section, we will compare psychophysical predictions of the model to behavioral data.

Based on neural and behavioral findings, we made a few simplifying assumptions that allowed the model dynamics to be solved for analytically (details in Methods). The sensory units are described by an *n*_*r*_-dimensional activity vector 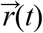. We assume that all of these sensory units receive the same time-varying input stimulus, *R*(*t*). Consistent with previous behavioral observations (Scott et al. 2015; Brunton, Botvinick, and Brody 2013; Pinto et al. 2018), our subjects’ performance did not depend on the duration of the pulse stream (Fig. 1g). We thus further assumed that the accumulators perform leak-free integration of their input; spatial dependence, to account for Fig. 1f, is described below. At each point in time, we took the input to the accumulator *a*_*r*_(*t*) to be a weighted sum of the activities of *n*_*r*_ sensory units. Thus 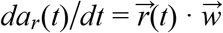, where 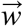 is the vector of input (feedforward) weights. A central feature of our model is that *a*_*r*_(*t*) feeds back as a dynamic, *multiplicative* gain control on the input to sensory units. By this we mean that the sensory responses 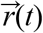 follow the input drive *R*(*t*) multiplied by a factor that is proportional to *a*_*r*_(*t*):

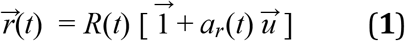

Here the vector 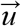 specifies the strength of the feedback from the accumulator to each sensory unit, and 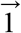 is the *n*_*r*_-dimensional vector with 1 for each coordinate. Note that, inspired by the brief transience of cue-locked neural responses (Fig. 2j), we have assumed in Eq. 1 that at each timepoint *t*, the sensory unit activities 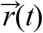 depend only on the current input *R*(*t*) and the current value of the accumulator *a*_*r*_(*t*). The feedback-loop model in Eq. 1 can produce either increasing or decreasing sensory responses *r*_*i*(*t*)_ vs. time, depending on whether the feedback weight *u*_*i*_ for the *i*^*th*^ sensory unit is positive or negative (illustrated in Fig. 5a).

**Figure 5.**
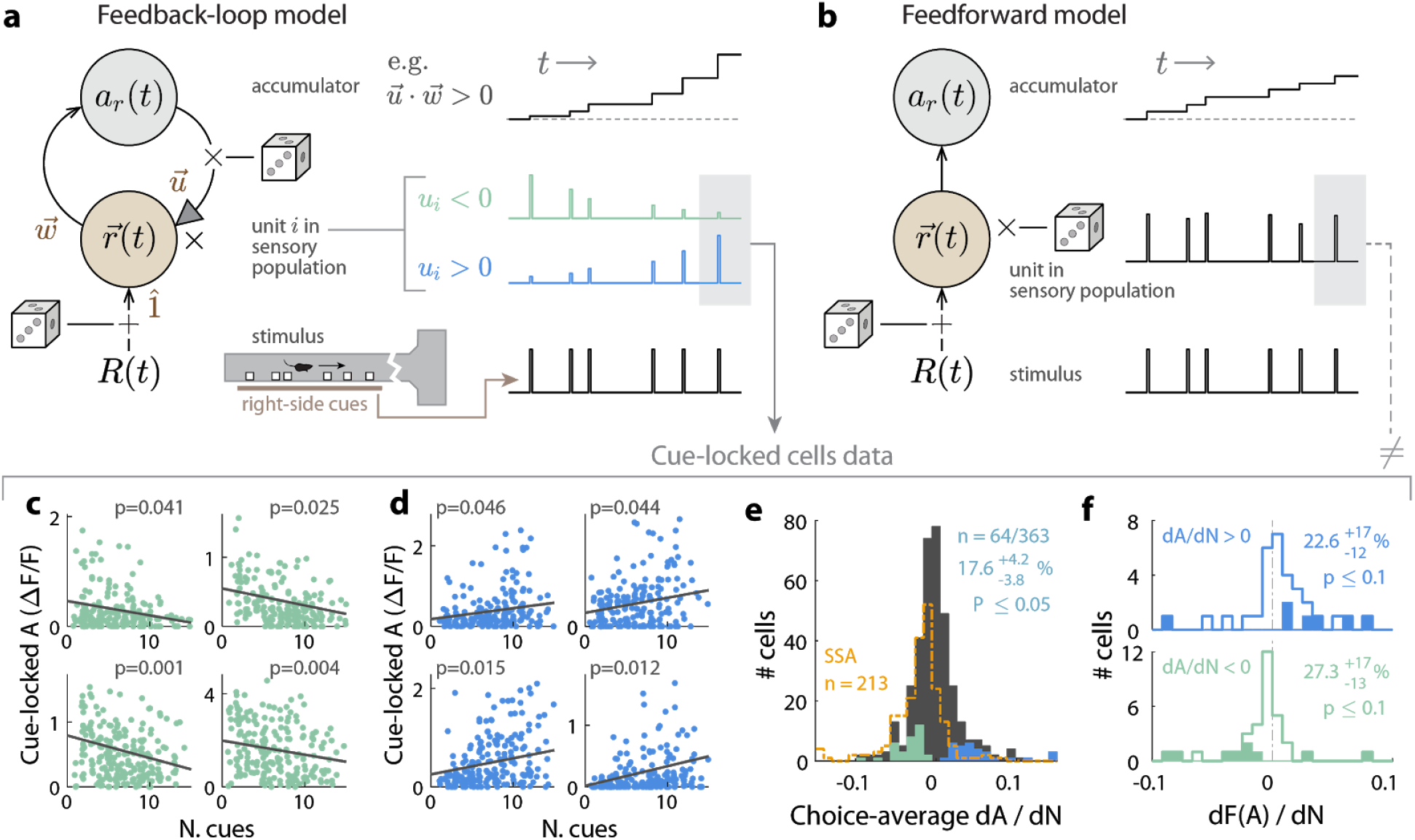
A multiplicative feedback-loop circuit can explain cue-locked cell amplitude modulations vs. cue counts. **(a)** Left: Schematic of the feedback-loop circuit model of a right-stimulus-stream accumulator. (Similar results and logic would apply to a left-stimulus-stream accumulator.) Dice indicate locations where additive/multiplicative noise may arise. Right: Single-trial illustration of time traces for dynamics at the sensory vs. accumulator stages of the model, for a given right-stimulus stream (bottom). When 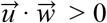, the accumulator state grows exponentially with the number of pulses. **(b)** As in (a), but for a purely feedforward model. The accumulator state grows linearly with the number of pulses. **(c)** Four example cue-locked cells (individual plots) with response amplitudes that were lower at the end of the cue region in trials with larger total numbers of cues (x-axis). Each point corresponds to the response of the cell to the last cue in a trial (gray shaded box in (a)). Lines are from a linear regression model that accounts for angular receptive field effects by weighting the data so that the φ_*cue*_ distribution is the same across cue-count bins. **(d)** As in (c), but for another four example cells with response amplitudes that were higher at the end of the cue region in trials with larger total numbers of cues. **(e)** Distribution of choice-averaged slope (*d*_*A*_/*d*_*N*_) for the linear regression model of amplitude *A* vs. number of preferred-side cues *N* as described in (c). Cells with significant slopes vs. a permutation test are highlighted in color (excluding cells compatible with SSA, dashed histogram). Numbers in the plot indicates the proportion of cells that had cue-count slopes that were significant at a *p* ≤ 0.05 level. **(f)** Distribution of slopes from linear regression of the Fano factor vs. preferred-side cue counts, separately for positively and negatively modulated cells as in (e). For cells with increasing (decreasing) amplitude dependence on cue counts, Fano factor tended to also increase (decrease) as a function of cue count, as predicted by the feedback model. Cells with significant Fano factor slopes vs. a permutation test are highlighted in color. Numbers in the plot indicate the proportion of cells with Fano factor slopes that were significant at a *p* ≤ 0.1 level.

Given the above, the accumulator dynamics are given by

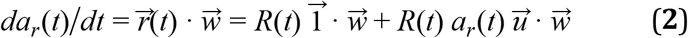

and the solution to this is (Methods):

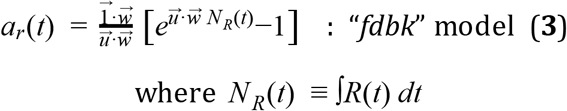

Because the accumulator output loops back to modulate sensory activity (Fig. 5a), and is thus again fed forward into the accumulator, the circuit can compound the effect of the stimulus and produce accumulator values that increase exponentially with the number of cues (Eq. 3). This is in contrast to a purely feedforward version, which would produce pure integration where the accumulator value *a*_*r*_(*t*) is proportional the the integral of the inputs *N*_*R*_(*t*) (obtained as a special case of Eq. 3 when 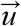 → 0):

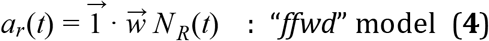

Note that the lack of a leak term in Eq. 3 and Eq. 4, and the multiplicative nature of the feedback in Eq. 2, means that when the input drive *R*(*t*) = 0 then *da*_*r*_(*t*)/*dt* = 0, i.e. the accumulators are stable. In fact, a feature of Eq. 3 and Eq. 4 is that the accumulator value does not depend explicitly on time, only on the cumulative stimulus input up to that time, *N*_*R*_(*t*) for arbitrary *R*(*t*). The intuition for pulsatile stimuli is that all changes to the sensory and accumulator states are gated by the stimulus drive, so the entire system only changes state when there is a pulse and is not sensitive to the time interval between pulses.

Consequently, psychophysical predictions and comparisons of these two models depend only on the net stimulus *N*_*R*_ at the end of a trial, and not otherwise on trial duration. In contrast to the multiplicative feedback we use here, it is also possible to achieve time-independent integration in an architecture with additive feedback, so long as there is also an appropriately tuned amount of accumulator leak (Seung 1996). However, in such circuits the accumulator continues to drive sensory units (defined as those that directly receive stimulus input) even in the absence of stimuli (Wimmer et al. 2015; Seung 1996). For pulsatile stimuli, this predicts changes in baseline sensory-unit activities in between pulses that would not match our observations of cue-locked cells (Fig. 2a-b,f), and we therefore did not consider additive feedback further. Instead, we focused on our multiplicative architecture, where sensory responses are scaled (Eq. 1) by an accumulator that depends on only the cumulative stimulus (Eq. 3), which is thus consistent with both neural and behavioral features that we set out to model (Fig. 1g, Fig. 2a-b,f). Although we do not have a normative explanation for the observed spatial weighting of evidence in the mouse behavioral data (Fig. 1f; not relevant for rats(Scott et al. 2015)), this can be empirically incorporated into the model by simply replacing the net stimulus *N*_*R*_ with a spatially-weighted sum of stimuli (see Methods and next section).

The feedback-loop model makes predictions about two properties of sensory unit activities that are compatible with the observed data: both pulse response amplitude and response variability should depend on cue counts. Before we probe these predictions in the neural data, we first point out that count-dependent stimulus response amplitudes could also be due to stimulus-specific adaptation (SSA). However, SSA, being inherently temporal, would then predict that behavioral performance decreases for shorter trial durations. In contrast, the behavior we examined did not depend on the interval between pulses (Fig. 1g, (Scott et al. 2015; Pinto et al. 2018); specifically, these task designs included a minimum interval between pulses to mitigate SSA effects). Consequently, for the following comparisons to neural data, we excluded cue-locked cells with responses consistent with SSA (Fig. 4c-d). Second, to minimize corrections for non-accumulator-related effects that could be due to other behavioral factors that depended on place/time in the trial (e.g. view angle), in the following analyses we restricted the data to responses to the last cue in the last third of the cue period, where there was little residual dependence of cue-locked responses on view angle and running speed (Supplementary Fig. 7a-b).

The first prediction of the feedback-loop model is that sensory units should respond to pulsatile stimuli with response amplitudes that are linearly related to the accumulator contents (Eq. 1). In the neural data, we thus asked whether the last-cue response amplitude *A* depended on the total number of cues *N* in that trial (i.e. cues of the preferred laterality for a given cell). We did this by linearly regressing *A* vs. *N*, controlling for choice by computing this separately in right- vs. left-choice trials, and controlling for view angle θ by weighting trials so that the θ distribution is the same across cue-counts for a fixed choice (Methods). Fig. 5c-d shows the *A* vs. *N* data for some example cells with choice-averaged *d*_*A*_/*d*_*N*_ slopes that were significant compared to a permutation test (Methods). These cells were part of two comparably sized subpopulations with significantly positive or negative choice-averaged *d*_*A*_/*d*_*N*_ slopes (Fig. 5e, cyan and green entries). In the model, these two subpopulations would correspond to positively-modulated (*u*_*i*_ > 0) and negatively-modulated *u*_*i*_ < 0) sensory units respectively. The second prediction of the feedback-loop model is that the Fano factor, *F* (*A*) ≡ *var*(*A*)/*mean*(*A*), should increase (decrease) with *N* for positively-(negatively-)modulated sensory units (Methods; illustrated in Supplementary Fig. 9a-b). Although other mechanisms can also produce non-constant Fano factors, a linear regression model of *F*(*A*) vs. *N* (Methods) indeed showed a compatible prevalence of positive (negative) Fano-factor slopes for the two subpopulations of cells with significantly positive (negative) *d*_*A*_/*d*_*N*_ slopes (Fig. 5f). The above count dependencies were also present in θ-controlled experiments (Supplementary Fig. 8; albeit statistical power was insufficient for the Fano factor analysis). In sum, the feedback-loop model predicts cue-count dependencies for amplitudes and variability of sensory unit activities that qualitatively match experimental observations for cue-locked cells. In contrast, purely feedforward architectures (Fig. 5b; see Eq. 4) predict sensory unit activities with no systematic count dependence.

### A multiplicative feedback-loop architecture best explains asymptotic Weber-Fechner scaling in psychophysical performance

Because a multiplicative feedback-loop circuit hypothesis can explain count-modulation of cue responses in the neural data, we wondered if it could also explain psychophysical features of pulsatile accumulation tasks. For otherwise identical models, the presence vs. absence of accumulator feedback on sensory units predicts two different mathematical forms (Eq. 3 vs. Eq. 4) for how left and right accumulator values depend on the input stimuli, and consequently potential differences in the accuracy of comparing two accumulators to produce a choice. We compared which of the feedforward versus feedback forms best fits two sets of behavioral data: the full Accumulating-Towers dataset (Pinto et al. 2018), as well as data from rats performing a visual pulse accumulation task with no navigational requirements (Scott et al. 2015). For all these data, the behavior had little dependence on time or trial history (Scott et al. 2015; Pinto et al. 2018), and the fraction of correct trials was only a function of the numbers of cues *N*_*maj*_ on the majority and *N*_*min*_ on the minority side. Two long-standing theories predict qualitatively different trends for this function, yet neither fully matches the data. As described below, our feedback model yields a better fit than either of these theories.

First, evidence accumulation is often modeled as a feedforward integration process, for which the central limit theorem of statistics states that performance should improve as a function of total pulse counts (Supplementary Fig. 9c, (Scott et al. 2015)). Alternatively, the psychophysical Weber-Fechner Law postulates that discriminability depends only on the ratio *N*_*min*_/*N*_*maj*_, i.e. performance should be constant as a function of total pulse counts *N*_*tot*_ = *N*_*maj*_ + *N*_*min*_ (Supplementary Fig. 9d, (Fechner 1966)). In both mouse and rat data (Fig. 6a), we intriguingly observed both types of trends. For *N*_*min*_/*N*_*maj*_ = 0 performance increased with *N*_*tot*_ as qualitatively predicted—but turns out *not* to be fully explained by—the central limit theorem and feedforward integration hypotheses. In contrast, for *N*_*min*_/*N*_*maj*_ = 0.4 there was little dependence of performance on *N*_*tot*_, consistent with the Weber-Fechner Law.

**Figure 6.**
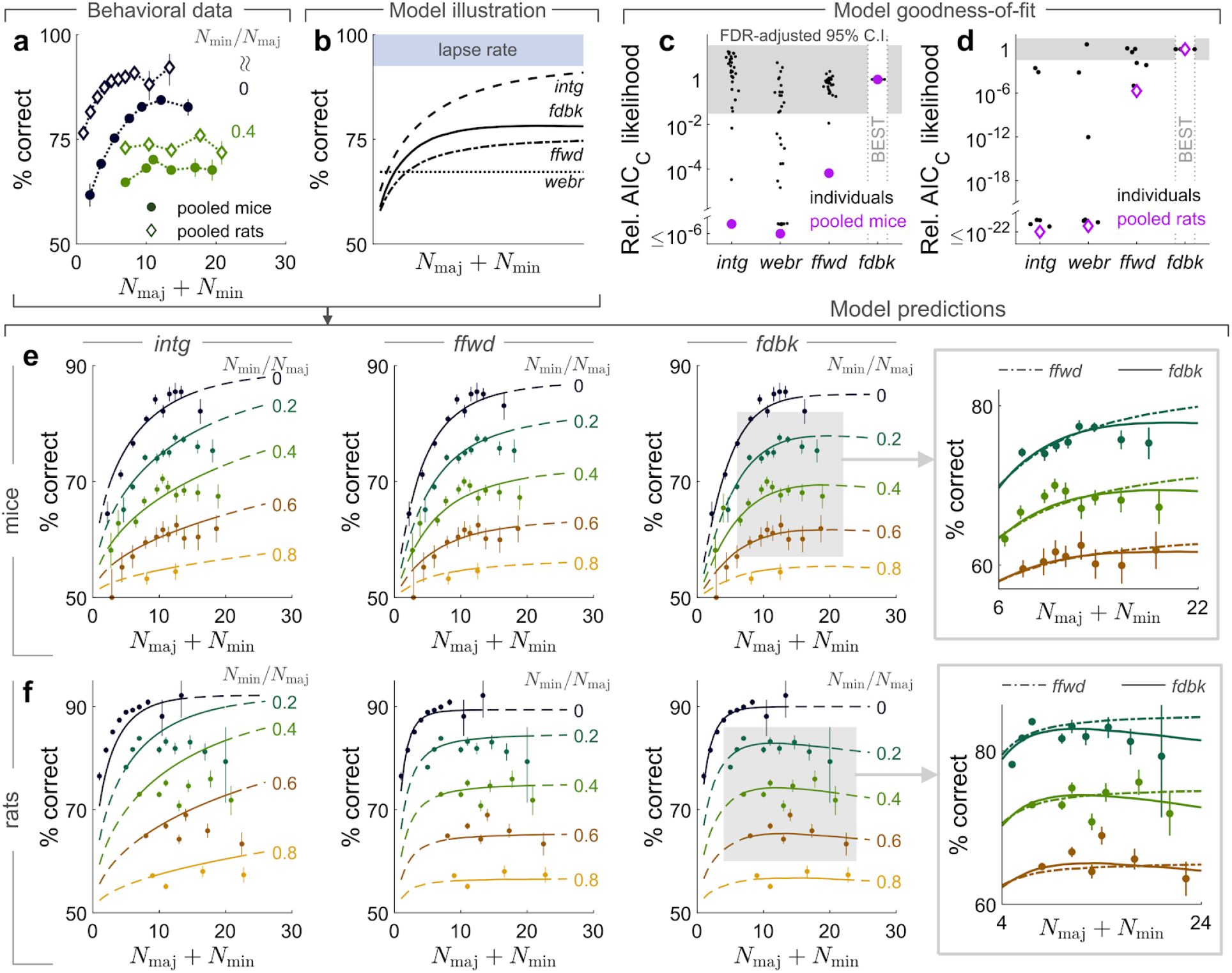
A multiplicative feedback-loop circuit best explains asymptotic Weber-Fechner scaling in perceptual performance, compared to models without feedback. **(a)** Behavioral performance (percent of correct trials) vs. total cue counts, for two fixed ratios of cue counts on the minority (*N*_*min*_) to majority (*N*_*maj*_) sides. Data were pooled across mice (rats). Points are joined by lines to guide the eye. Error bars: 68% C.I. across trials. **(b)** Illustration of perceptual discrimination accuracy for circuit architectures described in the text, with *N*_*min*_/*N*_*maj*_ = 0.5 and the same parameters for all noise distributions where relevant (σ1 = 1, μ*m* = 0, σ_*m*_ = 1, σ_*c*_ = 0.5, *p*_*lapse*_ = 0.15; see Methods for definition). The *webr* model predicts constant performance (parameters μ*m* = 0.1, σ_*m*_ = 1.6). **(c)** AIC_C_ likelihood ratios for models relative to the *fdbk* model, which best fits the pooled mouse data. Gray band: p-value threshold below which model goodness-of-fit are not significantly different from that of the best model, at an α = 0.05 test level after correcting for multiple comparisons using the Benjamini-Hochberg procedure (Benjamini and Hochberg 1995). **(d)** As in (c), but for rat behavioral data. **(e)** Pooled mouse data as in (d), compared to model predictions (lines) for three of the models illustrated in (b). The *webr* model (not shown) predicts constant performance vs. total counts. Error bars: 68% C.I. across trials. Rightmost box: zoomed-in view of data vs. *ffwd* (dash-dotted line) and *fdbk* (solid line) model predictions. **(f)** As in (e), but for pooled rat data vs. model predictions.

The above theories each assume that one dominant source of stochasticity drives the psychophysical performance of the accumulator circuit. For an integrator subject to the central limit theorem, this source is a per-pulse sensory noise that is independent across pulses. For pure Weber-Fechner scaling to apply, independent per-pulse sensory noise should be negligible compared to a memory-level noise (Gallistel and Gelman 2000). To account for the data deviating from both these theories, we hypothesized that the accumulator circuit instead mixes effects from multiple significant sources of stochasticity. In total, we considered the effects of (1) per-pulse sensory noise; (2) slow modulatory noise that multiplies the sensory response; and (3) noise associated with comparing accumulators (which has no count dependence). The critical comparison here is between two models that each had all of these three sources of stochasticity, but differed in whether there was accumulator-feedback onto the sensory units (“*fdbk*” model, Eq. 3), or if the circuit was purely feedforward (“*ffwd*” model, Eq. 4).

The framework that we used to evaluate the above models on behavioral data hypothesizes that the subject makes a decision by comparing the values of a right-side (*a*_*r*_) to a left-side (*a*_*l*_) accumulator, i.e. computing *a*_*r*_ − *a*_*l*_. In a given trial, the accumulator states *a*_*r*_ and *a*_*l*_ are each a sample from a distribution of possible (i.e. stochastic) values. How these distributions depend on the true pulse counts differentiates various accumulator architectures, as we will explain next. In addition, the operation of comparing two accumulators may itself be noisy, which we model as the subject forming the decision variable *c* = *a*_*r*_−*a*_*l*_ + δ where δ is a sample from another Gaussian distribution (“comparison noise”, with distribution variance 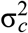). This decision variable *c* predicts that the subject will make a choice to the right with probability *P*(*c* > 0), up to an evidence-independent fraction *p*_*lapse*_ of lapse trials where the subject instead makes a random choice.

Here we summarize the definition of the *fdbk* and *ffwd* models, with details in the Methods. In both these models, the effect of sensory noise on the (say, right-side) accumulator output *a*_*r*_ in a given trial is to replace the true integrated stimulus *N*_*R*_ in Eq. 3 and Eq. 4 with a sample from a Gaussian distribution centered around *N*_*R*_, where the width of this distribution parameterizes the amount of sensory noise. Both models also have a modulatory noise source that randomly scales the amplitudes of sensory responses in a given trial. The difference between the two models is in where this modulatory noise arises in the hypothesized neural circuits. For the *fdbk* model we hypothesized that the accumulator feedback is noisy, or equivalently that the net feedback-loop gain (dot product of feedforward and feedback weights, 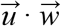 in Eq. 3) is stochastic per trial (Fig. 5a). This is in contrast to the *ffwd* model (where by definition 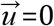), for which we hypothesized a non-specific source of slow gain fluctuations that scales the sensory responses by a stochastic value per trial (Fig. 5b).

The critical difference across these two models in how noise in *a*_*r*_ scales with the true integrated stimulus *N*_*r*_. We can understand this via an analysis of the variance of the *a*_*r*_ distribution. The simplest case is for the *ffwd* model, where including noise in Eq. 4 as discussed above yields 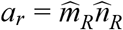, where 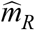 is a random modulatory noise variable with mean μ_*m*_ and variance 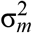, and 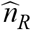 is a noisy integral of the sensory stimulus that has mean *N*_*R*_ and variance 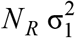 for a per-pulse noise variance 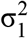 (central limit theorem). The variance of the *ffwd* accumulator output is thus:

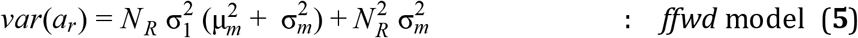

The first term in Eq. 5 corresponds to integration noise 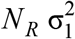 that is enhanced by modulatory noise effects 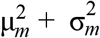, but as *N*_*R*_ increases it gradually becomes unimportant as the 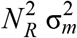 term dominates. This second term corresponds to Weber-Fechner scaling of perceptual uncertainty at high stimulus counts *N*_*R*_, i.e. the standard deviation of *a*_*r*_ becomes linearly proportional to *N*_*R*_. The hypothesized noise in comparing two accumulators adds a count-independent variance 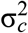 to the decision variable *c* = *a*_*r*_ − *a*_*l*_. Fig. 6b illustrates the effect on perceptual discrimination of these noise dependencies on *N*_*R*_. Here we fixed the ratio *N*_*min*_/*N*_*maj*_ = 0.5 of minority (*N*_*min*_ = the smaller of *N*_*R*_ or *N*_*L*_) to majority (*N*_*maj*_ = the larger of *N*_*R*_ or *N*_*L*_) side counts, and showed perceptual accuracy as a function of total cue counts *N*_*tot*_ = *N*_*maj*_ + *N*_*min*_. For ease of comparison to previously proposed theories, we explicitly consider two special cases of the *ffwd* model where all but one type of noise variance are set to zero: the integrator model which only has sensory noise (“*intg*” model, dashed line in Fig. 6b), and the Weber-Fechner model which only has modulatory noise (“*webr*” model, dotted line in Fig. 6b). At low total counts, the *ffwd* model performance (dash-dotted line in Fig. 6b) rises vs. *N*_*tot*_ as the effects of sensory and accumulator-comparison noises diminishes (cf. the *intg* model), but at high total counts the *ffwd* model performance saturates at less than the lapse rate because modulatory noise dominates (cf. the *webr* model).

Repeating the same exercise for the *fdbk* accumulator output, the noisy version of Eq. 3 is 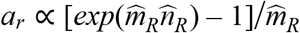 (up to an irrelevant constant factor), where 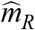 and 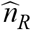 are stochastic variables same as in the *ffwd* model. A Taylor’s series expansion gives 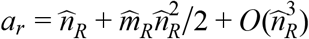, which has variance:

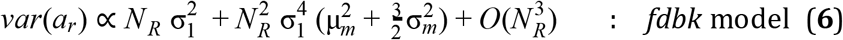

At low counts (*N*_*R*_ → 0) the *fdbk* model reduces to simple integration, which differs from the *ffwd* accumulator variance (Eq. 5) in that there is *no* scaling of the integration noise term 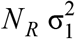 by modulatory noise effects. Intuitively, this is because at low counts (*N*_*R*_ → 0), the accumulator value also goes to *a*_*r*_ → 0, and therefore there is no gain on the modulatory noise. This means that for identical noise parameters, at low total counts the fdbk model performance (solid line in Fig. 6b) more closely follows the *intg* model performance rise vs. *N*_*tot*_ than the *ffwd* model. However at higher counts, the 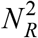 and higher-order terms in Eq. 6 can (for a large range of noise parameters) more quickly dominate over integration noise than for the *ffwd* model. In sum, the fdbk model can predict a steeper rise and an earlier saturation in performance vs. *Ntot* than the *ffwd* model (Fig. 6b). At asymptotically high counts, we show formally in the Methods that both model predictions converge and exhibit Weber-Fechner scaling.

The four accumulator models discussed above (*ffwd, fdbk,* and the two special cases of the *ffwd* model, the *intg* and *webr* models) have free parameters that specify the distributions of various noise sources. We estimated these for each model by maximizing model likelihoods with respect to the behavioral data (Methods). To allow for non-uniform spatial weighting of evidence in the mouse data (see Fig. 1f), when fitting these models to the behavioral data we used counts *N*_*R*_ that were spatially-weighted versions of the actual counts. The spatial weights were free parameters of the models, also fit to the data (Methods). The resulting best-fit spatial weights in the models showed a small primacy effect, as expected (Supplementary Fig. 10b, compared to Fig. 1f). Importantly, this spatial weighting had little dependence on the accumulator model architecture (Supplementary Fig. 10c) and thus did not affect our conclusions contrasting *ffwd* and *fdbk* models. For both mouse and rat data, the *fdbk* fit parameters indicated that the feedback-loop gain 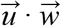 had distributions that were concentrated near zero (Supplementary Fig. 10d-f, the modes of the distributions were 0.018 for pooled mouse data and 0.006 for pooled rat data). This is compatible with our observations of near-equal proportions of positively (*u*_*i*_ > 0) vs. negatively (*u*_*i*_ < 0) count-modulated cue-locked cells in the mouse neural data (Fig. 5e), which can cancel each other out such that 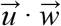 is typically small (see Discussion). Nevertheless, the trial-to-trial variability in 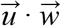 (corresponding to the long tail of the 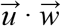 distribution) can predict appreciable effects on psychophysical performance trends.

The *fdbk* model best fitted the pooled data for both mice (Fig. 6c) and rats (Fig. 6d; see also residual distributions in Supplementary Fig. 10g-h). The behavior was particularly poorly explained by exclusively integration noise or Weber-Fechner scaling (*intg* and *wbr* models), instead strongly favoring models that involve a mixture of noise effects. We were unable to conclusively distinguish between models for most individual mice due to small sample sizes (particularly in parameter regions of high *N*_*tot*_), but Supplementary Fig. 10i shows significant trends of model likelihoods favoring the *fdbk* model for mice with greater number of trials. The larger rat datasets allowed identifying 3 out of 6 individual animals as significantly favoring the *fdbk* model over the *ffwd* model. No individual animal of either species favored the *ffwd* model over the *fdbk* model.

As discussed, the difference between the *ffwd* and *fdbk* models is in how quickly Weber-Fechner scaling starts to dominate with increasing *N*_*tot*_. Fig. 6e shows the pooled mouse data compared to predictions of three models. The *fdbk* model differs from the *ffwd* model in predicting a more constant asymptotic trend (see Supplementary Fig. 11a for individual mice). For rats, the *fdbk* model matches a visibly non-monotonic performance trend (Fig. 6f; Supplementary Fig. 11b for individual rats), which is mathematically impossible for the *intg* and *webr* models, and difficult to achieve in the *ffwd* model while simultaneously maintaining a good fit of the low-*N*_*tot*_ trend.

In sum, a circuit architecture with feedback modulations of sensory responses thus best explains rodent behavioral data. In addition, with respect to our mouse neural data, the *fdbk* model is the only model considered here that explains count dependence of cue-locked response amplitudes.

## Discussion

Psychophysics-motivated evidence accumulation models (Ratcliff and McKoon 2008; Stone 1960; Bogacz et al. 2006) have long guided research into how such algorithms may map onto neural activity and areas in the brain. A complementary, bottom-up approach starts from data-driven observations and formulates hypotheses based on the structure of the observations (Shadlen et al. 1996; Wimmer et al. 2015). In this direction, we exploited the mouse model system to systematically record from layers 2/3 and 5 of six posterior cortical areas during a task involving temporal accumulation of pulsatile visual evidence. A separate optogenetic perturbation study showed that all of these areas contributed to mice’s performance of the Accumulating-Towers task (Pinto et al. 2019). We reasoned that to understand how cortical areas contribute to evidence accumulation, a necessary first step is to understand the neural representation of sensory *inputs* to the process. In this work, we therefore focused on cue-locked cells that had sensory-like responses i.e. time-locked to individual pulses of evidence, which comprised ~5-10% of active neurons in visual areas and ~2% in the retrosplenial cortex (RSC). These cells are candidates for sensory inputs that may feed into an accumulation process that drives behavior, but could also reflect more complex neural dynamics such as from top-down feedback throughout the seconds-long decision formation process. We characterized properties of cue-locked responses across the posterior cortex, then asked if these properties suggest specific psychophysical effects in pulsatile evidence accumulation tasks.

One long-standing postulated function of the visual cortical hierarchy is to generate invariant visual representations (DiCarlo, Zoccolan, and Rust 2012), e.g. for the visual cues regardless of viewing perspective or placement in the T-maze. On the other hand, predictive processing theories propose that visual processing intricately incorporates multiple external and internal contextual information, in a continuous loop of hypothesis formation and checking (Rao and Ballard 1999; Bastos et al. 2012; Keller and Mrsic-Flogel 2018). Compatible with the latter hypotheses, we observed that across posterior cortices, cue-locked cells had amplitude modulations that reflected not only visual perspective and running speed (Niell and Stryker 2010; Saleem et al. 2013), but also the accumulated evidence, choice, and reward history (neural population decoding in Fig. 3). Inter-area differences were mostly in degree (Minderer, Brown, and Harvey 2019), with V1 having significantly lower performance for decoding view angle and choice, whereas RSC had lower decoding performance for speed but higher decoding performance for evidence (Fig. 3f). We also observed an anatomical progression from V1 to secondary visual areas to RSC in terms of increasing timescales of cue-locked responses (Fig. 2i-j) and increasing strengths of stimulus-specific adaptation (Fig. 4d). Our results are compatible with other experimental findings of increasing timescales along a cortical hierarchy (Murray et al. 2014; Runyan et al. 2017; Dotson et al. 2018; Schmolesky et al. 1998), and theoretical proposals that all cortical circuits contribute to accumulation with intrinsic timescales that follow a progression across brain areas (Hasson, Chen, and Honey 2015; Chaudhuri et al. 2015; Christophel et al. 2017; Sreenivasan, Vytlacil, and D’Esposito 2014).

The amplitude modulations of cue-locked cells can be interpreted as multiplicative gain changes on otherwise sensory responses, and could be clearly distinguished from additive effects due to our experimental design with pulsatile stimuli and high signal-to-noise calcium imaging (Fig. 2). While a number of other studies have quantified the presence of multiplicative noise correlations in cortical responses (Goris, Movshon, and Simoncelli 2014; Arandia-Romero et al. 2016; Lin et al. 2015), we showed that for most cells the amplitude variations were not random, but instead depended systematically on visuomotor and cognitive variables (Fig. 4c). Our findings extend previous reports of relationships between sensory responses and perceptual decisions, termed “choice probability (Britten et al. 1996)” (CP), and may constitute a form of conjunctive coding of cue and contextual information that preserves both the specificity and precise timing of responses to cues. An interesting question arises as to whether such multiplexing of cue and contextual information can cause potential interference between the different multiplexed information. For example, many evidence accumulation studies have reported positive correlations between CP and the stimulus selectivity of cells (Britten et al. 1996; Celebrini and Newsome 1994; Cohen and Newsome 2009; Dodd et al. 2001; Law and Gold 2009; Price and Born 2010; Kumano, Suda, and Uka 2016; Sasaki and Uka 2009; Gu, Angelaki, and DeAngelis 2014; Nienborg and Cumming 2014) (for a differing view, see (Zhao et al., n.d.) for a recent re-analysis at the neural-population level). Translated to our task, positive CP means that neurons that responded selectively to right cues tended to have increased firing rates when the animal will make a choice to the right. In this kind of coding scheme, increased activity in right-cue-locked cells could be due to either more right-side cues being presented or an internally generated right-choice signal, and there is no obvious way to distinguish between these two possibilities from just the activities of these cells.

Our data deviates from the abovementioned CP studies in that highly contralateral-cue-selective neurons could be divided into two near-equally sized subpopulations with positive choice modulation (analogous to CP > 0.5) and negative choice modulation (CP < 0.5) respectively (Supplementary Fig. 5e). In a closely related analysis, there were near-equal numbers of cue-locked cells with amplitudes positively vs. negatively modulated by accumulated counts (Fig. 5e). As two simultaneously recorded cells that respond to the *same* visual cue can be *oppositely* modulated (Fig. 4a), these phenomena are not expected from canonical accounts of spatial- or feature/object-based attention in visual processing (Cohen and Maunsell 2014; Treue 2014), but rather more compatible with mixed choice- and sensory-selectivity reported in other perceptual decision-making experiments (Raposo, Kaufman, and Churchland 2014). If choice- and count-modulations of cue response amplitudes originate from some form of accumulator feedback, our proposed feedback-loop (*fdbk*) circuit model suggests an interesting possibility. Intuitively, if comparable proportions of sensory units are positively vs. negatively modulated by feedback, the opposite signs of these modulations can cancel out when sensory unit activities are summed as input to the accumulator. In the model, 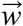 are feedforward weights (“synapses”) of sensory units onto the accumulator, and 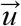 are feedback weights of the accumulator onto sensory units. The condition for cancellation is 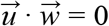, which reduces the accumulator dynamics (Eq. 2) to simple integration: 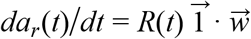. Therefore even with strong feedback connections (|*u*_*i*_| ≫ 0), the feedback-loop circuit can still be functionally equivalent to pure integration. We speculate that such a distribution of positive and negative feedback strengths across individual neurons in a population could be useful as a population-level “knob” for manipulating the *effective* gain of sensory input, e.g. feeding into processes such as evidence accumulation. The possibility for modulatory effects to cancel out is not limited to our feedback-loop model, as it applies generally to any readout of sensory unit activities. Similar arguments have been made for how motor preparatory activity and feedback do not interfere with motor output (Kaufman et al. 2014; Stavisky et al. 2017), how attentional-state signals can be distinguished from visual stimulus information (Snyder, Yu, and Smith 2018), and may hint at a general coding principle that allows non-destructive multiplexing of information in the same neuronal population.

Relationships between sensory responses and choice can arise in a purely feedforward circuit structure (Shadlen et al. 1996), where the causal role of sensory neurons in producing the behavioral choice predicts that choice-related neural and behavioral fluctuations should have similar timecourses (Nienborg and Cumming 2009). Incompatible with a solely feedforward circuit hypothesis, we instead observed that choice modulations of cue-locked responses *increased* in time (Fig. 4e) whereas the behavioral influence of sensory evidence fluctuations on the mice’s choice *decreased* in time (Fig. 1f). In a related analysis, we found that the response amplitudes of a substantial fraction of cue-locked cells actually depended on total cue counts in a graded manner (Fig. 5c-f). Both the choice- and count-modulation observations discussed here were suggestive of signals originating from an accumulator. Several features of our neural and behavioral data ruled out alternative possibilities such as stimulus-specific adaptation (details in Results).

Our neural observations above led us to propose the *fdbk* circuit model, where feedback from an accumulator acted as a dynamic multiplicative gain on sensory responses, and thus qualitatively explained how the response amplitudes of cue-locked cells could be modulated by the accumulated count of cues. Our motivation is similar in spirit to (Wimmer et al. 2015), which investigated a related neural network model where neurons in a sensory circuit were recurrently coupled to neurons in an integration circuit via feedforward as well as (additive) feedback connections. These authors showed that the feedback connections in their model simulations were required to qualitatively explain the experimentally observed opposite timecourses of choice-related neural and behavioral fluctuations in (Nienborg and Cumming 2009). A key difference with respect to our *fdbk* model is that here the accumulator feedback onto sensory responses is multiplicative, as opposed to additive, as this better matches our neural observations of transient, amplitude-modulated sensory responses (Fig. 2a-b,f) as well as behavioral observations that performance does not depend on pulse stream duration (Fig. 1g). A main focus of our work is also to quantitatively compare hypotheses of accumulator architectures with and without feedback (*fdbk* vs. *ffwd* models), in terms of how well they can predict psychophysical trends in rodent behavioral data. As we discuss next, we found that the data clearly favored the *fdbk* model. In this way, our work connects neural to behavioral observations via a suggested neural circuit mechanism.

From a methodological standpoint, we used a simple dynamical systems model for the *fdbk* vs. *ffwd* circuits that allowed us to analytically examine the impact of different sources of noise that could arise in the components of these circuits. In particular, we aimed to understand how the accuracy of discriminating whether there were more right- or left-side counts depended on total pulse counts for a fixed ratio of counts on either side. Two incompatible predictions of this dependence are made by two famous theories. One, if accumulation is an integration process, then the central limit theorem of statistics predicts that performance should increase with total counts. Two, the long-standing Weber-Fechner law of perception postulates that there should be no change in performance vs. total counts (Fechner 1966). In contrast to these theories, the *fdbk* model hypothesizes that multiple significant sources of noise are at play: (1) noise in comparing two accumulators, which has a constant scale and quickly becomes insignificant vs. total counts; (2) independent noise in the detection of each sensory pulse, which has gradually diminishing effect with total counts; and (3) noise in the accumulator feedback gain on sensory responses, the effect of which does not change with total counts and therefore produces asymptotic Weber-Fechner scaling. Both the mouse and rat behavioral data showed deviations from Weber-Fechner law but only at low total counts, and these trends could be much better explained by our *fdbk* model than either of the aforementioned famous theories (Fig. 6c-d). The *fdbk* model also better explained pooled mouse and rat data than the *ffwd* model, which had all the same sources of stochasticity except that (3) was replaced by random gain fluctuations per trial (unrelated to accumulator values, since this model has no feedback), equivalent to a memory-level noise as previously hypothesized to cause Weber-Fechner scaling (Gallistel and Gelman 2000). For both rat and mouse data, the optimal parameters for the psychophysical *fdbk* model indicated that the net gain around the feedback loop (dot product of feedforward and feedback weights 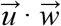) was small (Supplementary Fig. 10d-f; compatible with the mouse neural data Fig. 5e). Nevertheless, because these trial-to-trial fluctuations in multiplicative feedback-loop gain are proportional to the signal magnitude, at high enough counts they dominate over the other sources of noise and produce Weber-Fechner scaling in perceptual accuracy.

We note that the *fdbk* model describes the effective structure of the computational circuit, but does not constrain anatomical details of where the computation takes place. The biological implementation of multiplicative feedback may be an input-output property of individual neurons (Peña and Konishi 2001; Silver 2010), which for pulsatile inputs may be a threshold-linear as opposed to truly multiplicative operation. Alternatively, multiplication can be a network computation, for example involving an intermediate population of inhibitory neurons (Olsen et al. 2012; Atallah et al. 2012; Wilson et al. 2012; Zhang et al. 2014; Fu et al. 2014; Pi et al. 2013; S. Lee et al. 2013; S.-H. Lee et al. 2012). While a multiplicative feedback-loop architecture results in suboptimal perceptual discrimination if compared to simple integration (Fig. 6b; assuming all else kept equal), it may reflect other behavioral and engineering pressures that brains have evolved to handle. For example, amplification may be important for selecting a tiny relevant signal out of a massive amount of sensory data regarding the world, and perhaps also out of a massive amount of other neural activity in the brain itself.

## Acknowledgements

We thank B.B. Scott for brainstorming and feedback on the concept of this paper, as well as L. Pinto, C.M. Constantinople, A.G. Bondy, M. Aoi, and B. Deverett for useful and interesting discussions. B. Engelhard and L. Pinto built rigs for the high-throughput training of mice, and S. Stein helped in the training of mice in this study. B. Engelhard and L. Pinto contributed behavioral data from the mouse evidence accumulation task. B.B. Scott and C.M. Constantinople contributed behavioral data from the rat evidence accumulation task. We additionally thank all members of the BRAIN COGS team, Tank and Brody labs. This work was supported by the NIH grants 5U01NS090541 and 1U19NS104648, and the Simons Collaboration on the Global Brain (SCGB).

## Author Contributions

SAK performed the experiments, data analysis and conceptualization/modeling. SYT and DWT designed the experimental setups. SAK wrote the manuscript with input from DWT and CDB. SAK, DWT, and CDB conceived the project.

## Declaration of Interests

The authors declare no competing interests.

## Methods

### Experimental Model and Subject Details

All procedures were approved by the Institutional Animal Care and Use Committee at Princeton University and were performed in accordance with the Guide for the Care and Use of Laboratory Animals(National Research Council et al. 2011). We used 11 mice for the main experiments (+4 mice for control experiments), aged 2-16 months of both genders, and from three transgenic strains (see Supplementary Table 2) that express the calcium-sensitive fluorescent indicator GCamp6f(Chen et al. 2013) in excitatory neurons of the neocortex:

- 6 (+2 control) mice (6 male, 2 female): Thy1-GCaMP6f(Dana et al. 2014) [C57BL/6J-Tg(Thy1-GCaMP6f)GP5.3Dkim/J, Jackson Laboratories, stock # 028280]. Abbreviated as “Thy1 GP5.3” mice.
- 5 (+1 control) mice (3 male, 3 female): Triple transgenic crosses expressing GCaMP6f under the CaMKII α promoter, from the following two lines: Ai93-D; CaMKII α-tTA [IgS5^tm93.1(tetO−GCaMP6f)Hze^ Tg(Camk2atTA) 1Mmay/J(Gorski et al. 2002), Jackson Laboratories, stock #024108] (Madisen et al. 2015); Emx1-IRES-Cre [B6.129S2-Emx1^tm1(cre)Krj^/J, Jackson Laboratories, stock #005628]. Abbreviated as “Ai93-Emx1” mice.
- 1 mouse (control experiments; female): quadruple transgenic cross expressing GCaMP6f in the cytoplasm and the mCherry protein in the nucleus. both Cre-dependent, from the three lines: Ai93-D; CaMKII α-tTA, Emx1-IRES-Cre, and Rosa26 LSL H2B mCherry [B6;129S-Gt(ROSA)26Sor^tm1.1Ksvo^/J, Jackson Laboratories, stock #023139].

Mice were randomly assigned such that there were about the same numbers of either gender and various transgenic lines in each group (main vs. control experiments). As the Ai93-Emx1 strain had higher expression levels of the fluorescent indicator, they produced significantly higher signal-to-noise (SNR) recordings than the Thy1 GP5.3 strain, and contributed more to the layer 5 datasets (see Supplementary Table 2). Strain differences in the results were small and not of a qualitative nature (Supplementary Fig. 1d-g, Supplementary Fig. 5h).

### General statistics

We summarize the distribution of a given quantity vs. areas and layers using quantile-based statistics, which are less sensitive to non-Gaussian tails. The standard deviation is computed as half the difference between the 84% and 16% quantiles of the data points. The standard error (S.E.M.) is computed as the standard deviation divided by 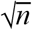 where *n* is the number of data points. For uncertainties on fractions/proportions, we compute a binomial confidence interval using a formulation with the equal-tailed Jeffreys prior interval (DasGupta, Tony Cai, and Brown 2001). The significance of differences in means of distributions were assessed using a two-sided Wilcoxon rank sum test. The p-value threshold for evaluating significance is 0.05 for all tests, unless otherwise stated.

### Behavioral metrics

These analyses were described in a previous study (Pinto et al. 2018) and outlined here. The fraction of trials where a given mouse turned right was computed in 11 bins of evidence levels Δ ≡ #*R* − #*L* at the end of each trial, and fit to a 4-parameter sigmoid function *pR*(Δ) = *p*_0_ + *B*[1 + *e*^−(Δ−Δ0)/λ^]^−1^ to obtain psychometric curves. A logistic regression model was used to assess the dependence of the mice’s choices on the spatial location of cues, i.e. with factors being the evidence {Δ_*i*_|*i* = 1, 2, 3} computed using cues in equally-sized thirds of the cue region (indexed by *i*). Statistical uncertainties on the regression weights were determined by repeating this fit using 1000 bootstrapped pseudo-experiments.

### Decoding from cue-locked amplitudes

The decoding models were fit separately using responses to cues in three equally sized spatial bins of the cue region. We defined the neural state response as the vector of contralateral-cue-locked cell response amplitudes to a given cue, and used a Support Vector Machine classifier (SVM) to predict a task variable of interest from this neural state (using data across trials but restricted to responses to cues in a given third of the cue region, as mentioned). To assess the performance of these classifiers using 3-fold cross-validation, we trained the SVM using 2/3rds of the data and computed Pearson’s correlation coefficient between the predicted and actual task variable values in the held-out 1/3rd of the data. Significance was assessed by constructing 100 null hypothesis pseudo-experiments where The Δ*F*/*F* of cells for a given epoch bin were permuted across trials.

To correct for multiple comparisons when determining whether the decoding p-value for a particular dataset was significant, we used the Benjamini-Hochberg procedure(Benjamini and Hochberg 1995) as follows. For a given type of decoder, we sorted the *p*-values of all data points (spatial bins and imaging sessions) in ascending order, [*p*_1_, *p*_2_,…, *p*_*n*_], and found the first rank *i*_α_ such that 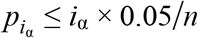. The decoding performance was then considered to be significantly above chance for all 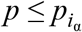.

### Uncorrelated modes of task variables

We wished to define a set of uncorrelated behavioral modes such that the original set of six task variables are each a linear combination of these modes, with the additional requirement that each mode should be as similar as possible to one of the task variables. In matrix notation, this means that we want to solve:

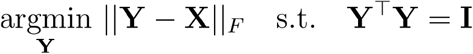

where each column of **X** corresponds to values of a given task variable across trials, each column of **Y** are the uncorrelated behavioral modes, and 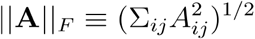 is the Frobenius norm of a matrix **A**. This can be computed using polar decomposition (Higham 1988): **X** = **YH**, where **Y** is an orthogonal matrix and **H** a symmetric matrix. To obtain the polar decomposition **X** = **UΣV**^Τ^, we used an algorithm based on the singular value decomposition, which gives the solution **Y** = **UV**^Τ^.

### Impulse response model for cue-locked cells

This analysis excluded some rare trials where the mouse backtracks through the T-maze, by using only trials where the *y* displacement between two consecutive behavioral iterations was > −0.2*cm* (including all time-points up to the entry to the T-maze arm), and if the duration of the trial up to and not including the ITI was no more than 50% different from the median trial duration in that session.

We modeled the activity of each cell as a time series of non-negative amplitudes *A*_*i*_ in response to the *i*^*th*^ cue, convolved with a parametric impulse response function *g*(*t*):

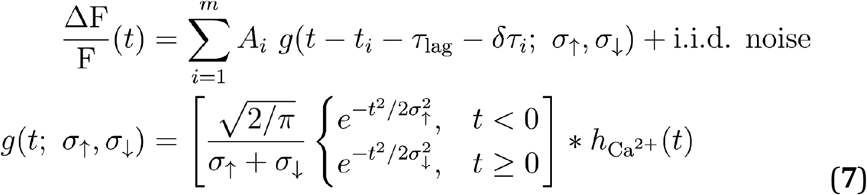

where {*t*_*i*_|*i* = 1,…, *m*} are the appearance times of cues throughout the behavioral session. The free parameters of this model are the lag (τ_*lag*_), rise (σ_↑_) and fall (σ_↓_) times of the impulse response function, the amplitudes *A*_*i*_, and small (L2-regularized) time jitters δτ_*i*_ that decorrelates variability in response timings from amplitude changes. 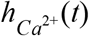 is a calcium indicator response function using parameters from literature (Chen et al. 2013), which deconvolves calcium and indicator dynamics from our reports of timescales. This function is parameterized as a difference of exponentials, 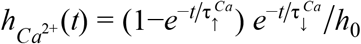, where 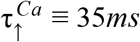, 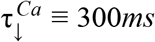 and *h*_0_ is a normalization constant such that the peak of this function is 1. We note that timescales cannot be resolved to better than about the imaging data rate of 1/15*Hz* ≈ 67*ms*.

We maximized the model likelihood to obtain point estimates of all the parameters, using a custom coordinate-descent-like algorithm (Wright 2015). The significance of a given cell’s time-locking to cues was defined as the number of standard deviations that the impulse response model AIC_C_ score (bias-corrected Aikaike Information Criterion (Hurvich and Tsai 1989)) lies above the median AIC_C_ of null hypothesis models where the timings {*t*_*i*_} of cues were randomly shuffled within the cue region. Given the Δ*F* /*F* time-series *F* (*t*) for a given cell and the predicted activity time-trace *m*(*t*), the AIC_C_ score is:

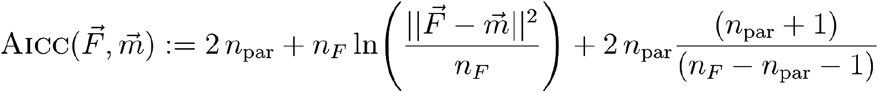

where *n*_*F*_ is the number of time-points that comprise the data and *n*_*par*_ is the number of free parameters in the model. Lastly, a small fraction of cells responded to both left- and right-side cues. We parsimoniously allowed for different impulse responses to these by first selecting a primary response (preferred-side cues) as that which yields the best single-side model AIC_C_, then adding a secondary response if and only if it would improve the fit. We defined cells to be cue-locked if the primary response significance exceeded 5 standard deviations.

### Amplitude modulation models

These models used as input the following behavioral data: *t*_*i*_ is the onset time of the *i*^*th*^ cue, which is located at distance *y*_*i*_ along the cue region, and appears at a visual angle relative to the mouse (Fig. 1c). Δ(*t*_*i*_) is the cumulative cue counts (explained further below) up to and including cue *i*, and *C*(*t*_*i*_) is the upcoming choice of the mouse in that trial. *v*_*i*_ ≡ *v*(*t*_*i*_) is the running speed of the mouse in the virtual world at the time that the *i*^th^ cue appeared, and for simple linear speed dependencies explained below, the standardized version 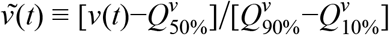 is used, where 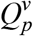 is the *p* probability content quantile of the speed distribution.

To account for the stochastic and nonnegative nature of pulsatile responses, the cue-locked cell response amplitudes *A*_*i*_ were modeled as random samples from a Gamma distribution, *A*_*i*_ | *μ*_*A*_(*t*_*i*_),*k* ~ Γ[*k*,*μ*_*A*_(*t*_*i*_/*k*]. The shape parameter *k* for the Gamma distribution is a free parameter, and furthermore indexed by choice for the choice model. The four models discussed in the text are defined by having different behavior-dependent mean functions **μ**_*A*_(*t*_*i*_) that have the following forms (detailed below):

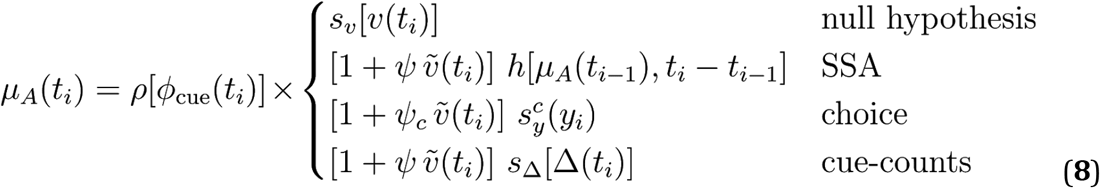

In all of the models, *ρ*(*ϕ*_cue_) is an angular receptive field function that has either a skew-Gaussian (Priebe, Lisberger, and Movshon 2006) or sigmoidal dependence on *ϕ*_cue_:

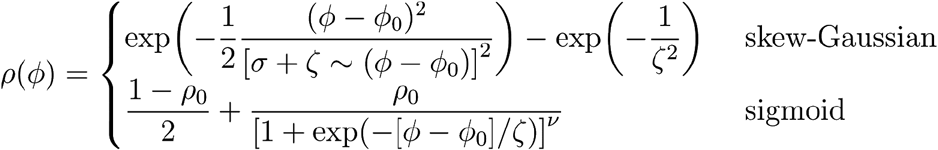

*ϕ*_0_,*σ*,*ζ*, *ρ*_0_ and *ν* are all free parameters, and either the skew-Gaussian or sigmoidal hypotheses are selected depending on which produces a better fit for the cell (using the AIC_C_ score as explained below).

All of the models also have a speed dependence that multiplies the angular receptive field function. For the null hypothesis, we allowed this to be highly flexible so as to potentially match the explanatory power of the other models (which have other behavioral dependencies). Specifically, the function *s*_*v*_(*v*) is defined to be a cubic spline (piecewise 3rd-order polynomial (Gan 2004)) with control points at five equally-spaced quantiles of the running speed distribution, i.e. at 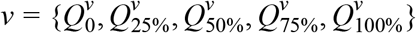. A cubic spline model has as many free parameters as the number of control points. For the other models, we used a simple linear parameterization for speed dependence, 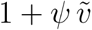 where *ψ* is a free parameter (for the choice model, there are two free parameters *ψ*_0_ where *C* indexes the choice).

The SSA, choice, and cue-counts models are further distinguished by how they depend on the *h*, 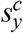, and *s*_Δ_ functions respectively. For the SSA model:

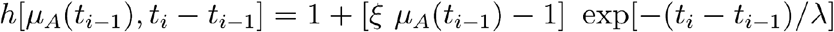

The response to the first cue in the session is defined to be μ_*A*_(*t*_1_) = 1. The *h* function can be understood as follows. Right after the cue at *t*_*i*−1_, the response is scaled by the free parameter ξ, i.e. the new response level is ξ μ_*A*_(*t*_*i*−1_) where ξ > 1 corresponds to facilitation and ξ < 1 corresponds to depression. This facilitation/depression effect decays exponentially with time towards 1, i.e. the amount by which the response μ_*A*_(*t*_*i*_) deviates from 1 is equal to the deviation (from 1) of the facilitated/depressed response ξ μ_*A*_(*t*_*i*−1_) − 1, multiplied by the time-recovery factor exp[−(*t*_*i*_ − *t*_*i*−1_/λ]. Here λ is another free parameter that specifies the timescale of recovery.

The choice model has smooth dependencies on *y* location on the cue region parameterized by choice. This is given by two functions 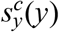 where *C* indexes either the right or left choice, and each of these functions is a cubic spline with control points at *y* = {0, 0.5*L*_*cue*_, *L*_*cue*_} (recall that *L*_*cue*_ is the total length of the cue region).

Lastly, the cue-counts model also has smooth dependencies on cue counts Δ, i.e. the function *s*_Δ_(Δ) is a cubic spline. As the responses of cells can depend on counts on either the right, left, or both sides (Scott et al. 2017), we allowed Δ to be either the cumulative right or cumulative left cue counts (control points are at Δ = {0, 3, 8}), or the cumulative difference #*R* − #*L* in cue counts (control points are at Δ = {− 4, 0, 4}). The best definition of Δ was selected per cell according to which produced the best AIC_C_ score.

Because neural activity can be very different in the rare cases where the mouse halts in the middle of the cue region, only data where the speed *v* is within 25% of its median value were included in the analysis of this model. Point estimates for the model parameters were obtained by minimizing the Gamma-distribution negative log-likelihood:

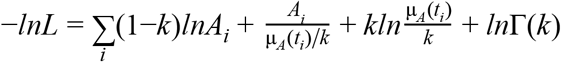

Because the Gamma distribution is defined only in the positive domain, we had to make an assumption about how to treat data points where *A*_i_ = 0. We reasoned that we could substitute these with a noise-like distribution of amplitudes, which were obtained by fitting the impulse response model (Eq. 7) using the same cue timings but simulated noise-only data, which comprised of a Δ*F*/*F* time-series drawn i.i.d. from a Gaussian distribution with zero mean and standard deviation being σ^*F*^, the estimated fluorescence noise level for that cell. The relative AIC_C_-based likelihood used for model selection as described in the text, is *exp*([*AIC*_*C*_(*model* 1)−*AIC*_*C*_(*model* 2)]/2).

### Choice modulation strength

The location-dependent choice modulation strength for cue-locked amplitudes is defined as 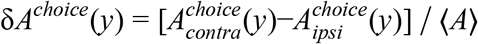, where 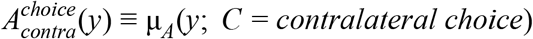 as in Eq. 8, and analogously for ipsilateral choices. This is computed by evaluating the amplitude model prediction vs. location in the cue region, but at fixed φ_*cue*_ corresponding to zero view angle (+ 22° for right-side cues and − 22° for left-side cues) and Δ = 0. The normalization constant is:

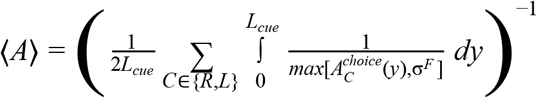

### Feedback-loop model solution

As explained in the main text, the sensory and accumulator states are specified by:

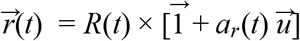

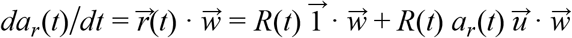

This is an ordinary differential equation that can be solved by the integrating factor method. Writing it in the form 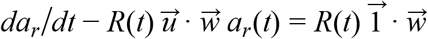, we identify the integrating factor to be 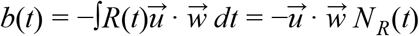, giving a solution:

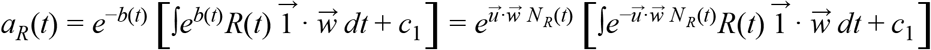

where *c*_1_ is a constant of the integration. This can be simplified to:

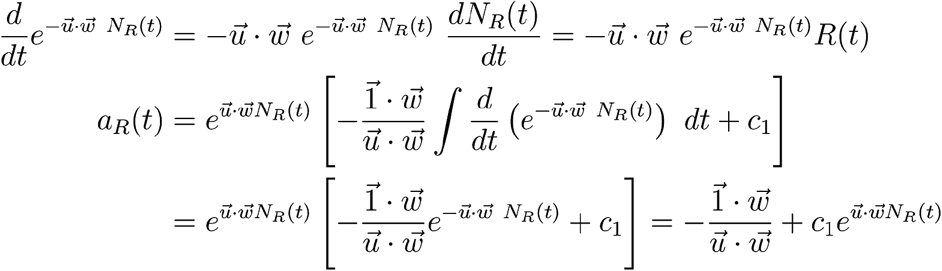

We assume that at the start of the trial, the accumulator has zero content. This requires 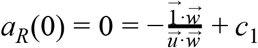, and substituting 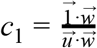 into the above we obtain Eq. 3.

In the case of weak feedback, a Taylor’s series expansion of Eq. 3 gives 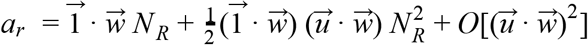 The accumulator thus reduces to perfect integration in the zero feedback limit (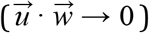), and exhibits growing nonlinearities vs. *N*_*R*_ for stronger feedback. The relevant measure of feedback strength is not the “synaptic” weights ^→^*u*, but the projection of the feedback onto the feedforward direction, 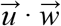. In other words, near-linear integration can be achieved in a neural circuit with strong feedback synapses, so long as there is net cancellation such that 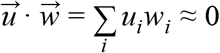. This requires the sensory population state to be at least 2-dimensional.

### Dependence of last-cue response amplitudes on counts

We fit a different linear model 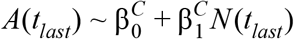 for each subset of trials with choice *C* ∈ {*right*, *lef t*}, where *A*(*t*_*last*_) is the amplitude of the response to the last cue in the last third of the cue period, and *N* (*t*_*last*_) is the total number of preferred-side cues in that trial. To control for visual angle (Runyan et al. 2017), we weighted the data used in this fit so that the φ*cue* distributions were the same in three equally sized quantile bins when conditioned on *N* ∈ {1 − 3, 4 − 6, 7 − 9, ≥ 10}. Similarly, the Fano factor *F* (*A*) ≡ *var*(*A*)/*mean*(*A*) was computed per *N*-bin as the weighted variance over weighted mean. Linear regression was then performed for *F* (*A*) vs. the bin-average *N*.

Null hypotheses were constructed by shuffling the amplitudes *A* across trials for a fixed choice *C*, and the slope 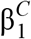 for a cell is considered to be significant if less than 5% of shuffled-data fits have slopes greater than that value. For cells with either significant 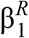 or 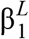, these slopes were highly significantly correlated across choice categories (Supplementary Fig. 7c), and we therefore used the choice-averaged slope 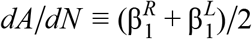 to identify significantly modulated cells. The significance of this choice-averaged slope was analogously defined by comparing its value to the choice-averaged slopes in shuffled-data fits.

### Predicted Fano factor of cue-locked responses

Here we consider the theoretical consequences of *N*_*R*_ in Eq. 3 not being perfectly deterministic, but instead Gaussian-distributed random variate with mean *N*_*R*_ and variance 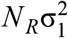 (as per the central limit theorem as explained in the text). This means that 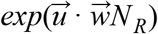 is lognormally distributed with non-logarithmized mean 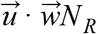 and variance 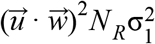. Letting 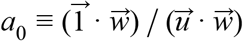, 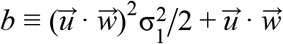 and 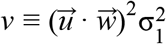to simplify notation, the accumulator state has the following mean and variance (Johnson, Kotz, and Balakrishnan 1994):

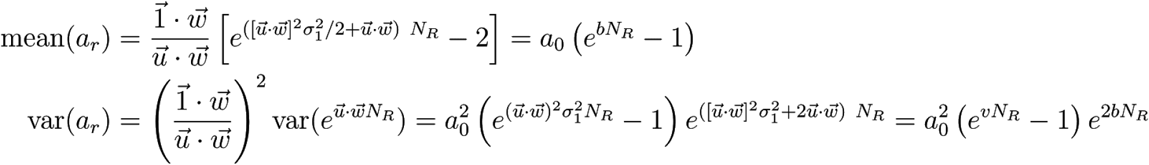

We want to understand the trend of the Fano factor vs. *N*_*R*_ for the activity of a single sensory neuron under pulsatile input. From Eq. 1 this is *ri* = *R* (1 + *a_r_ui*) where *a*_*r*_ is the stochastic accumulator state above and *R* corresponding to a single input pulse is a Gaussian random variate with mean 1 and variance 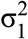. For simplicity we will assume that *R* and *a*_*r*_ are independent random variables, which would be the case in the discrete-time version of Eq. 3 since for causality reasons the scaling by the accumulator state should depend on the inputs up to but not including *R*. We use the following rule for the variance of a product of two independent random variables:

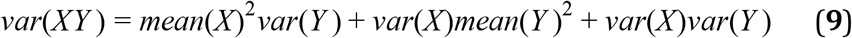

The Fano factor of the distribution of *r*_*i*_ is thus:

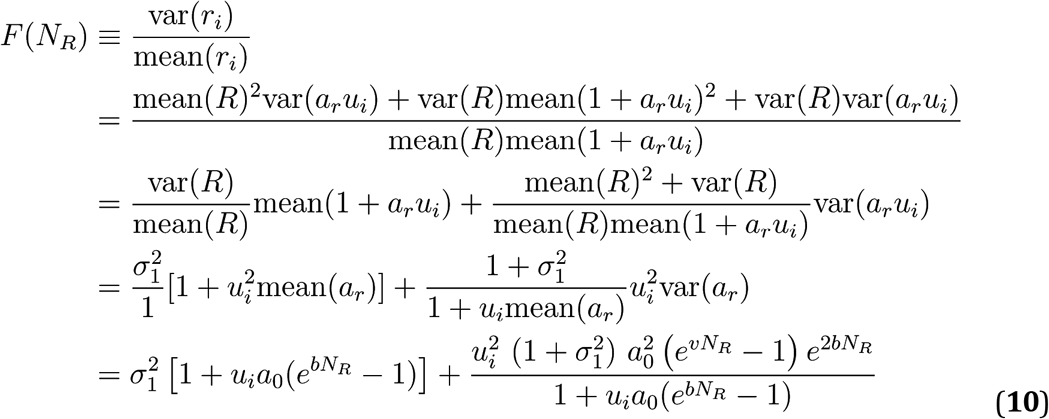

In the regime of small *N*_*R*_ we expand Eq. 10 in a Taylor’s series, 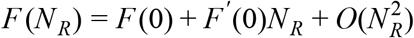. Direct substitution of *N*_*R*_ = 0 in the above gives us 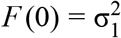. For the first-order term, we have:

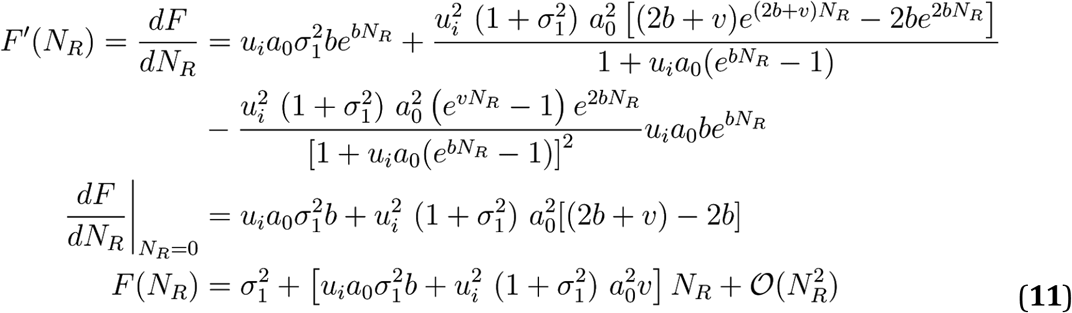

All the variables *a*_0_, *b*, *v*, and σ_1_ are positive, whereas *u*_*i*_ can take on either sign. This means that for cells with feedback strength *u*_*i*_ > 0, all the terms in Eq. 11 are non-negative, in particular the first-order term a.k.a. the slope, and *F* is thus a monotonically increasing function of *N*_*R*_. Viewed as a function of the feedback strength, *f* (*u*_*i*_) ≡ *F* ^′^(0) has roots at 0 and 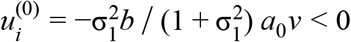, so for *u*_*i*_ < 0 cells there is a range of 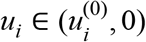 where *F*′ (0) < 0 i.e. *F* is a monotonically decreasing function of *N*_*R*_, for small enough *N*_*R*_ such that the Eq. 11 expansion holds. However as *N*_*R*_ grows large but still for *u*_*i*_ < 0, this trend reverses. We can see this by considering the behavior of *F* (*N*_*R*_) close to the singular point where *mean*(1 + *a_r_u_i_*) → 0. Going back to the form of Eq. 10 as 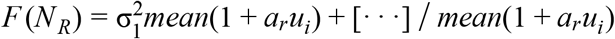, the first term proportional to the mean becomes negligible compared to the second which is inversely proportional. The latter diverges to + ∞, because the denominator decreases monotonically from 1 down to 0, while the numerator is always non-negative. The latter follows from the squares of real numbers always being non-negative, which is also the case for those of the form *e^x^* ≥ 1 when all *x* ≥ 0. Thus to recap, for *u*_*i*_ < 0 cells the Fano factor first decreases with *N*_*R*_, then at some point turns to increasing since it diverges to + ∞ as *mean*(1 + *a_r_u_i_*) → 0. For extremely strong negative feedback such that the mean sensory response would decrease below zero, we must extend the model to include rectification effects to prevent this from happening. This is beyond the scope of this work, although depending on the type of rectification, this can also modify the behavior of the Fano factor (Charles et al. 2018).

We illustrate the above analytical results using simulations where for a given true input count of *N*, we model the single-pulse stochasticity *R* as a Gamma-distributed random variable with mean 1 and variance σ^2^_1_ the accumulator state as Eq. 3 but with *N*_*R*_ replaced with a Gamma-distributed random variable with mean *N* −1 and variance 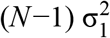 (discrete-time version). The parameters used are 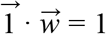, 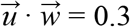 and *u*_*i*_ =± 0.01. The resulting distribution of sensory responses as in Eq. 1 is shown in Supplementary Fig. 9a, but bounded to be no less than zero. For *N* = 1, the accumulator state is zero and thus there is no distinction between positively and negatively modulated sensory responses. For *N* = 5 and *N* = 10 the feedback effect grows progressively larger, inducing a separation between *u*_*i*_ > 0 and *u*_*i*_ < 0 cases. Supplementary Fig. 9b shows the mean, variance, and Fano factor of the sensory response distributions for a range of true counts. The Fano factor monotonically increases for *u*_*i*_ > 0 and has the expected non-monotonic trend of first decreasing then increasing for the *u*_*i*_ < 0 case given our choice of parameters. These results are not sensitive to the type of distribution (e.g. Gamma vs. Gaussian) modeled, so long as the variance remains finite and there is not an excessive amount of probability mass that is truncated at zero.

### Sources of noise in various accumulator architectures

To account for imperfect psychophysical performance, we consider three sources of noise in modeling the distribution of *a*_*r*_.

First is sensory noise, any source of stochasticity that occurs either in the stimulus *R*(*t*) itself or in the activity state 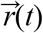 of the sensory units. Since only the projection of the sensory state onto the readout direction 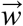 matters in terms of changes to the accumulator state, only the net 1-dimensional variability along 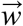 matters. Sensory noise is thus equivalent to replacing *R*(*t*) → *R*(*t*) + ε(*t*) where ε(*t*) is a (scalar) noise process. Assuming the central limit theorem, we model this as replacing the true integral 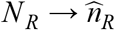 where 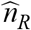 is a Gaussian distributed random variable with mean *N*_*R*_ and spread 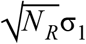.

Secondly, we consider modulatory noise that may be present in gain modulations of the sensory units. For the *fdbk* model, as the accumulator modulation of sensory responses is proportional to the feedback weight, this is equivalent to there being stochasticity in the feedback weights 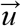, which for tractability we consider to be slow on the timescale of a trial. Again, only the 1-dimensional variability along 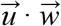 matters, so in sum this source of stochasticity leaves Eq. 3 unchanged but instead can be considered as 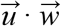 being drawn randomly per trial, 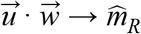. We model 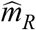 with a lognormal distribution, which is the limiting case of a product of many positive random variables. For the *ffwd* model, we instead hypothesized a non-specific source of slow gain fluctuations, equivalent to scaling all sensory unit responses with a stochastic variable per trial, 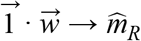 in Eq. 4, where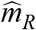 has a lognormal distribution.

Third, for both models there can be noise associated with comparing the two accumulators to form a decision, which we model as the decision variable *c* being Gaussian-distributed around *a*_*r*_−*a*_*l*_.

Sources of variability that for simplicity we did not model include initial and drift noise in the accumulator, as these were found to be negligible according to behavioral models of pulsatile evidence accumulation (Brunton, Botvinick, and Brody 2013; Scott et al. 2015). Accumulator drift noise also predicts a time-dependence to the behavior that we do not observe. However, note that for the *ffwd* model, initial noise in the accumulator would be interchangeable with comparison noise in the decision variable *c*.

Formally, the modified differential equation for the evolution of the *fdbk* model accumulator state given a sensory noise process ∊(*t*) and a modulatory noise process ξ(*t*) is:

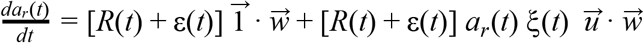

The solution as discussed above (assuming that ξ(*t*) is slower than the trial duration) is:

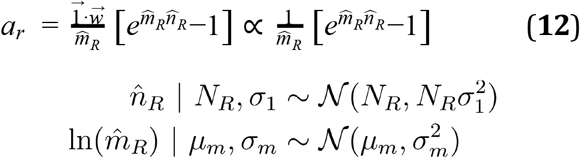

The model predicts that the animal should make a choice to the right with probability given by the decision variable 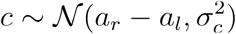 being positive. The likelihood of getting exactly (*k*_*R*_, *k*_*L*_) right- and left-choice trials in the behavioral data for a fixed (*N*_*R*_, *N*_*L*_) is binomially distributed around this probability. The constant 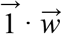 that *a*_*r*_ is proportional to can be neglected since this just sets the arbitrary units of the accumulator, which we assume that the brain adjusts so that the right and left accumulators can be compared in a way that is not biased by having potentially different units. We additionally modeled a lapse rate that acts as the animal instead making a fair coin toss to go right *p*_*lapse*_ of the time, instead of the evidence-dependent probabilities above. Lastly, we maximized the model likelihood over the five free parameters {σ_1_, μ_*m*_, σ_*m*_, σ_*c*_, *p_lapse_*} to best fit the behavioral data as described in the next section.

As discussed in the text, we assessed how well the *fdbk* model predicts behavior in comparison to three other accumulator architectures. The *ffwd* model has accumulator states distributed as 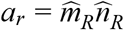, and the full set of five free parameters {σ_1_, μ_*m*_, σ_*m*_, σ_*c*_, *p_lapse_*}, same as for the *fdbk* model. The *intg* variant of this model has no modulatory nor accumulator-comparison noise (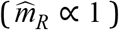), and so has two free parameters {σ_1_, *p*_*lapse*_}. The *webr* variant instead has no sensory nor accumulator-comparison noise 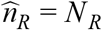, and so has three free parameters {μ_*m*_, σ_*m*_, *p*_*lapse*_}. The modulatory noise 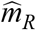 is the cause of Weber-Fechner scaling in all three of the *webr*, *fdbk* and *ffwd* models.

### Fitting accumulator models to behavioral data

The behavioral accumulator models used data from 17 mice (144366 trials from all mice with ≥ 500 trials) that performed the Accumulating-Towers task [Citation error], and data from 6 rats (266984 trials) that performed a head-fixed flash-accumulation task (Scott et al. 2015).

Let *k*_*R*_(*N*_*R*_, *N*_*L*_) and *k*_*L*_(*N*_*R*_, *N*_*L*_) be the number of right- and left-choice trials respectively out of all trials in the behavioral data with a given number of true right (*N*_*R*_) and left (*N*_*L*_) cue counts. A particular accumulator model *M* ∈ {*intg*, *webr*, *ffwd*, *fdbk*} specifies a probability of right-choice trials given the true cue counts, 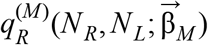 where 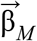 are the free parameters of the model. Including a lapse rate *p*_*lapse*_, we optimized the models by minimizing their negative log-likelihood, which have the same form as given by the binomial probability distribution:

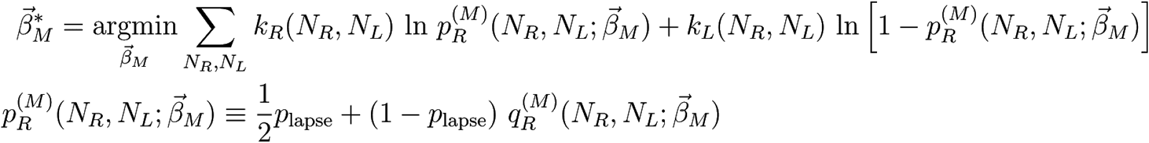

For the *intg* model, the probability distribution of right- and left-accumulator states are Gaussian according to the central limit theorem. Assuming that the accumulators are independent and have the same parameters, the joint probability distribution is 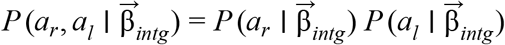, and the right-choice probability is given by the region 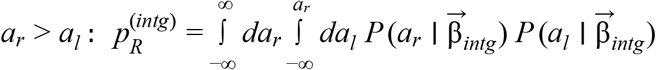. This has a well-known simplification in that the difference *c* ≡ *a*_*r*_−*a*_*l*_ of two (independent) Gaussian-distributed variables is also Gaussian-distributed, so 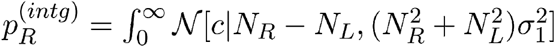.

For the remaining three models, we evaluated the necessary integrals using a Monte Carlo method. For a given combination of (*N_R_*, *N_L_*), we drew 200,000 samples each for the five stochastic quantities 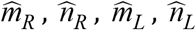 (see Eq. 12), and 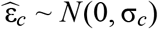 which acts as the accumulator comparison noise. The decision variable was computed as 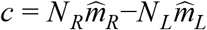 for the *webr* model, 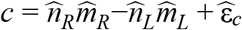 for the *ffwd* model, and 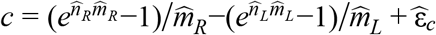 for the *fdbk* model. The right-choice probabilities 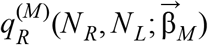 for these model were then estimated as the fraction out of the 200,000 samples for which *c* > 0.

For the mouse data, we included spatial weights for evidence as free parameters in all models (cf. Fig. 1f). This means that instead of the models having free parameters 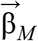 described above, they have parameters 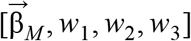 where *w*_1_,…, *w*_3_ are the evidence-weights for three equally sized spatial bins of the cue region. For each trial, we computed the number of right-side cues in these spatial bins: 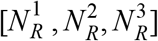 such that 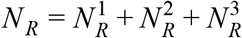, and analogously for left-side cues. The right-choice probability for each model was then computed as 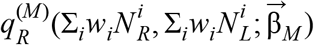, i.e. with 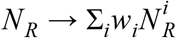 being the spatially-weighted number of right-side cues. Because an overall scaling of *w*_*i*_ is fully redundant with the noise scales already in all accumulator models, we imposed the constraint *w*_1_ + *w*_2_ + *w*_3_ = 3 as part of the model fitting procedure. These parameters should therefore be interpreted as relative weights for evidence in the three spatial bins.

Interestingly, for the mouse *fdbk* model fits, accumulator-comparison noise had a more dominant effect than sensory noise (Supplementary Fig. 10a-left), whereas the reverse holds for rats (Supplementary Fig. 10b-left). This could be related to the stimuli in the rat task being very brief (10ms) LED flashes, compared to the high visual salience of cues in the mouse task. The navigational aspect of the mouse task instead likely adds unmodeled sources of behavioral variability that may have been absorbed into the accumulator-comparison noise term.

### Asymptotic Weber-Fechner scaling of feedback-loop model performance

Even though Eq. 3 predicts that the *fdbk* model accumulator state is exponentially related to the true counts, this exponential scale does not on its own predict any change in perceptual performance if there is only sensory noise. This is because if 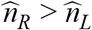 for the perfect integrator, then 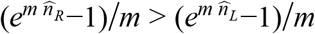 since we have only applied the same monotonic transformation to both left- and right-hand-sides (assuming *m* > 0). What distinguishes the *ffwd* and *fdbk* models is therefore the interaction of sensory and other sources of noise. In particular, modulatory noise is required for both models to exhibit Weber-Fechner scaling in perceptual discrimination.

We analytically show that the *fdbk* model makes the same predictions as the *ffwd* model at asymptotically high pulse counts, as follows. As previously explained, the noisy *fdbk* accumulator state is 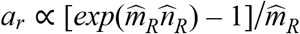 for a stochastic modulatory variable 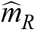. At high counts 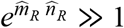, so 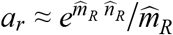. Making use of the fact that *ln a*_*r*_ > *ln a_l_* if and only if *a*_*r*_ > *a*_*l*_, we can equivalently consider the performance of comparing the log-transform of the accumulator output, 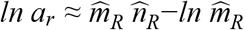. Already 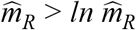, so at high counts *N_R_* → ∞ where 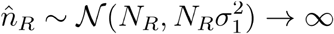, we quickly have 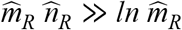 and therefore 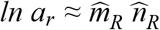. To summarize, comparing two *fdbk* accumulators is equivalent to comparing 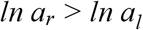, and at high counts this becomes approximately the same as comparing 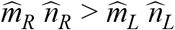 (which is precisely the *ffwd* model Eq. 4). The *fdbk* model thus asymptotically converges to the *ffwd* model, and in the same way, Weber-Fechner scaling.

## Supplemental Information

**Supplementary Table 1.**
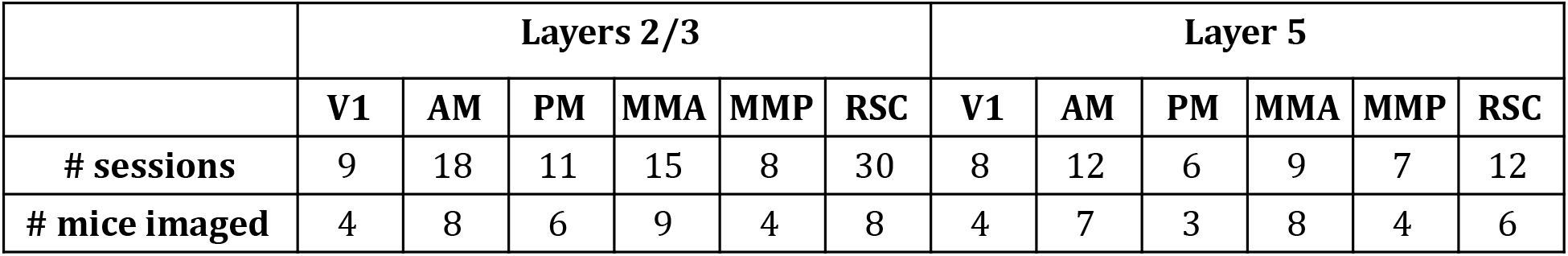
Number of imaging sessions and mice for various areas and layers, for the main experiment.

|  | Layers 2/3 |  |  |  |  |  | Layer 5 |  |  |  |  |  |
| --- | --- | --- | --- | --- | --- | --- | --- | --- | --- | --- | --- | --- |
|  | V1 | AM | PM | MMA | MMP | RSC | V1 | AM | PM | MMA | MMP | RSC |
| # sessions | 9 | 18 | 11 | 15 | 8 | 30 | 8 | 12 | 6 | 9 | 7 | 12 |
| # mice imaged | 4 | 8 | 6 | 9 | 4 | 8 | 4 | 7 | 3 | 8 | 4 | 6 |

**Supplementary Table 2.** Overall performance and number of imaging sessions for the main experiment, per mouse (rows), in various areas and layers (columns). Mice of the Thy1 GP5.3 strain have names starting with “gp”, and those from the Ai93-Emx1 strain have names starting with “ai” (see Methods).

|  |  | Layers 2/3 |  |  |  |  |  | Layer 5 |  |  |  |  |  |
| --- | --- | --- | --- | --- | --- | --- | --- | --- | --- | --- | --- | --- | --- |
|  | % correct | V1 | AM | PM | MMA | MMP | RSC | V1 | AM | PM | MMA | MMP | RSC |
| gp31 | 73.0 | 1 | 1 | 1 | 1 | 0 | 2 | 1 | 1 | 1 | 1 | 1 | 1 |
| gp36 | 75.3 | 2 | 1 | 1 | 1 | 1 | 2 | 2 | 1 | 1 | 1 | 1 | 1 |
| gp37 | 70.1 | 0 | 1 | 0 | 1 | 1 | 1 | 0 | 0 | 0 | 1 | 0 | 0 |
| gp40 | 67.7 | 0 | 2 | 0 | 0 | 0 | 9 | 0 | 1 | 0 | 0 | 0 | 5 |
| gp42 | 67.1 | 0 | 1 | 0 | 2 | 3 | 4 | 0 | 1 | 0 | 1 | 2 | 1 |
| gp46 | 67.8 | 0 | 0 | 1 | 3 | 0 | 4 | 0 | 0 | 0 | 1 | 0 | 2 |
| ai50 | 68.7 | 0 | 2 | 1 | 3 | 0 | 7 | 0 | 1 | 0 | 2 | 0 | 2 |
| ai53 | 67.8 | 2 | 0 | 0 | 1 | 0 | 0 | 2 | 0 | 0 | 0 | 0 | 0 |
| ai55 | 71.6 | 0 | 3 | 0 | 2 | 0 | 1 | 0 | 3 | 0 | 1 | 0 | 0 |
| ai56 | 71.9 | 4 | 7 | 6 | 1 | 3 | 0 | 3 | 4 | 4 | 1 | 3 | 0 |
| ai57 | 67.2 | 0 | 0 | 1 | 0 | 0 | 0 | 0 | 0 | 0 | 0 | 0 | 0 |

**Supplementary Figure 1.**
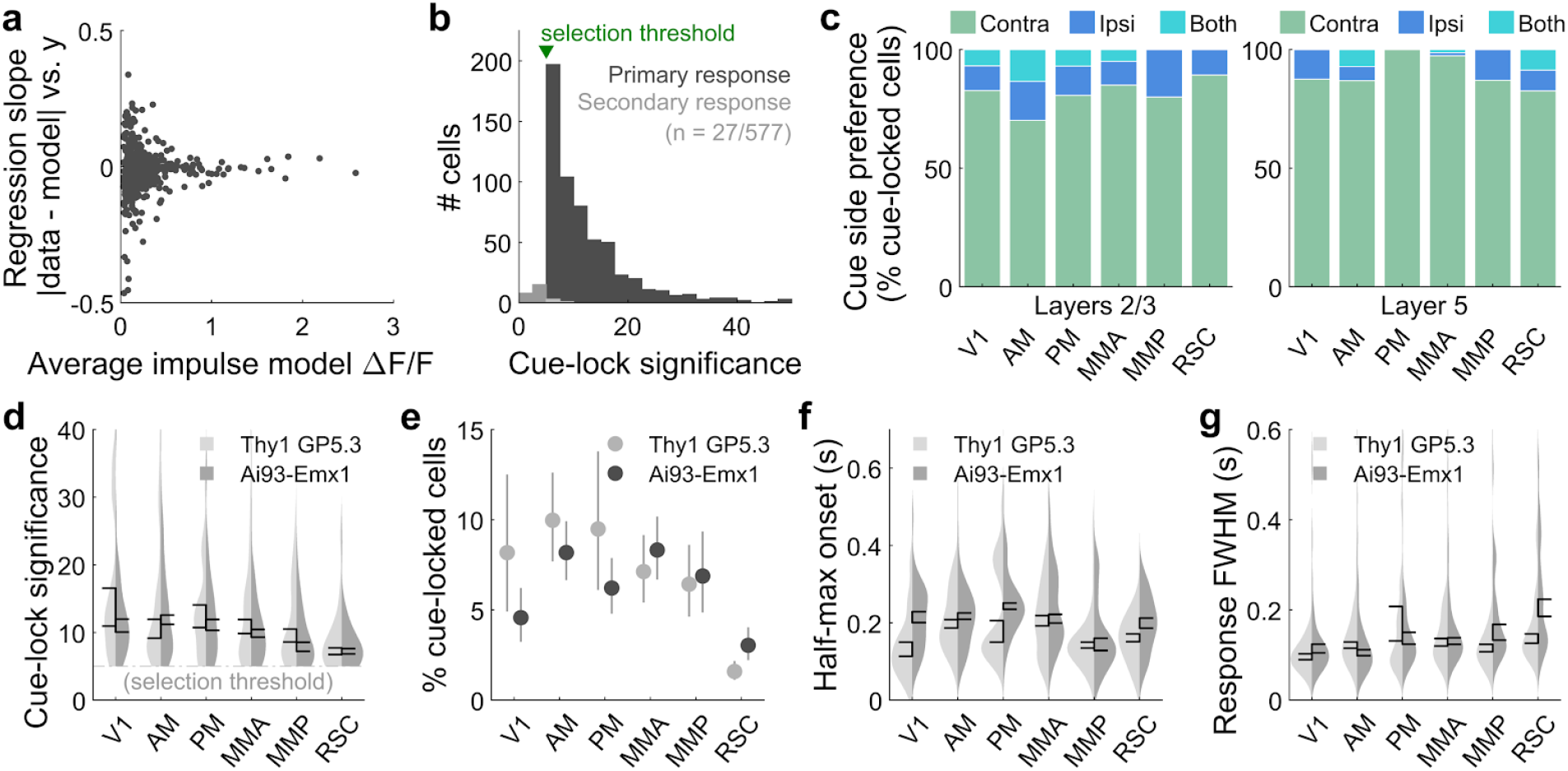
Additional statistics for cue-locked responses. **(a)** Slope (β_1_) from a linear regression model ρ = β_0_ + β_1_*y*, where ρ is the residual magnitude (absolute value of data minus impulse response model prediction) shown in Fig. 2f, and *y* is the spatial bin of the cue region in which the residual was computed (as explained for Fig. 2f). Each point corresponds to a single significantly cue-locked cell. Residuals were normalized per cell by the average impulse model prediction (across trials and place in the cue region) for that cell; this normalization factor is also shown as the x-coordinate in the plot. Units were defined for the spatial bins such that 0 (1) corresponds to the start (end) of the cue region. A slope of ± 1 can therefore be interpreted as a change in residuals from the start to the end of the cue region by an amount comparable to the mean signal predicted by the impulse response model. **(b)** Distribution of cue-locking significance for cells with a significant primary response (above 5 standard deviations compared to cues-shuffled fits). **(c)** Proportion of cells in various areas/layers that respond only to contralateral cues (green), only ipsilateral cues (dark blue), or to cues on both sides (light blue). **(d-g)** As in Fig. 2g-j, but comparing data from two strains of mice. Data were pooled across layers.

**Supplementary Figure 2.**
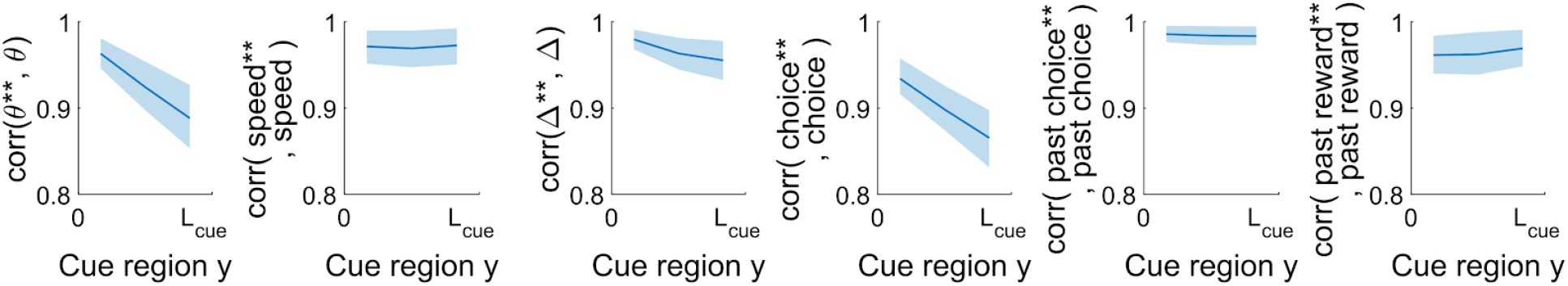
Pearson’s correlation between uncorrelated behavioral modes (θ **, speed**, etc.) and the corresponding most similar task variable (θ, speed, etc.). These uncorrelated modes were computed via polar decomposition as explained in the Methods. Bands: standard deviation across imaging sessions.

**Supplementary Figure 3.**
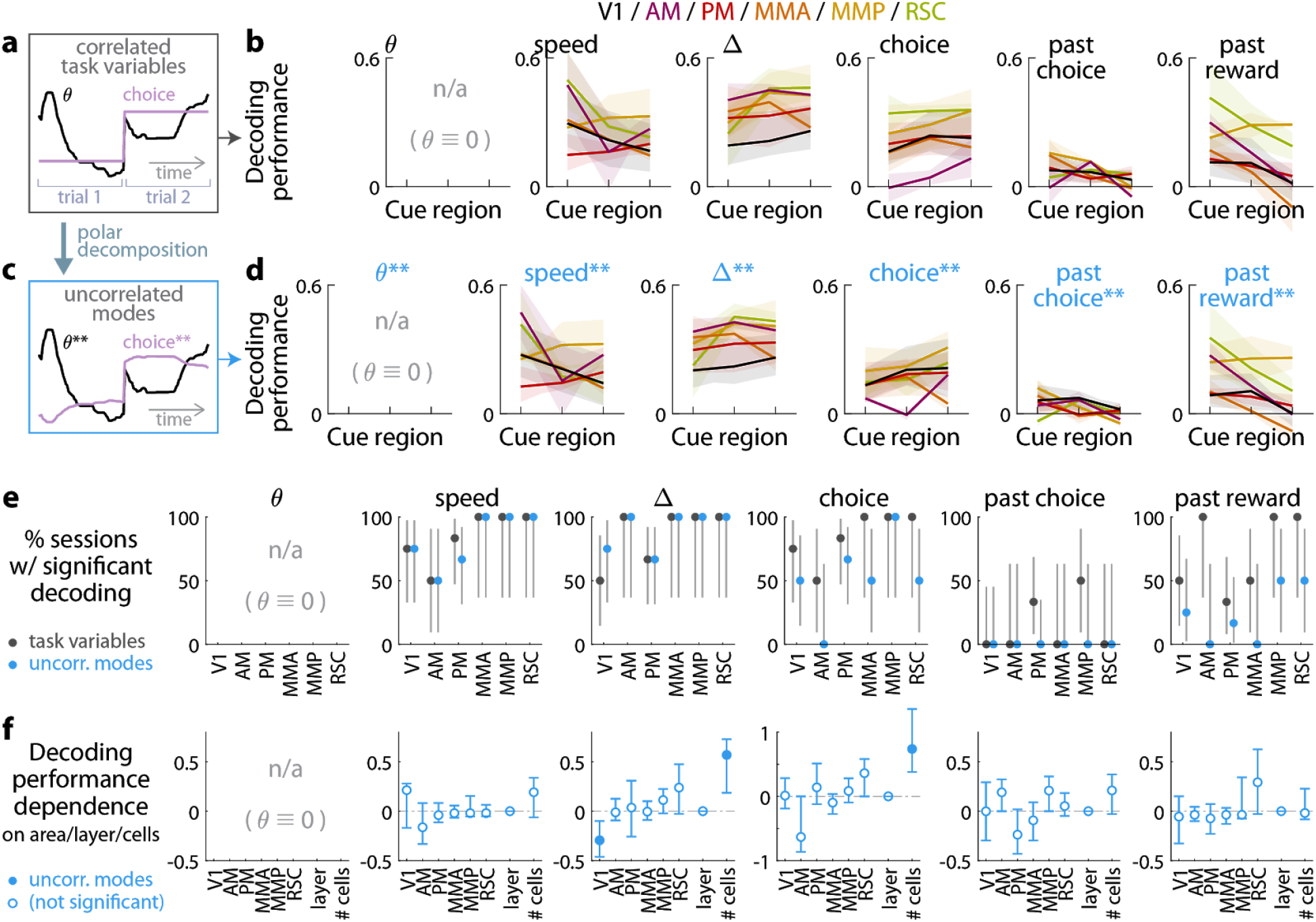
Qualitatively similar performances for decoding task variables from cue-locked response amplitudes in control experiments with view-angle restricted to zero in the cue region. (a-f) As in Fig. 3, but using data from the θ-controlled experiments.

**Supplementary Figure 4.**
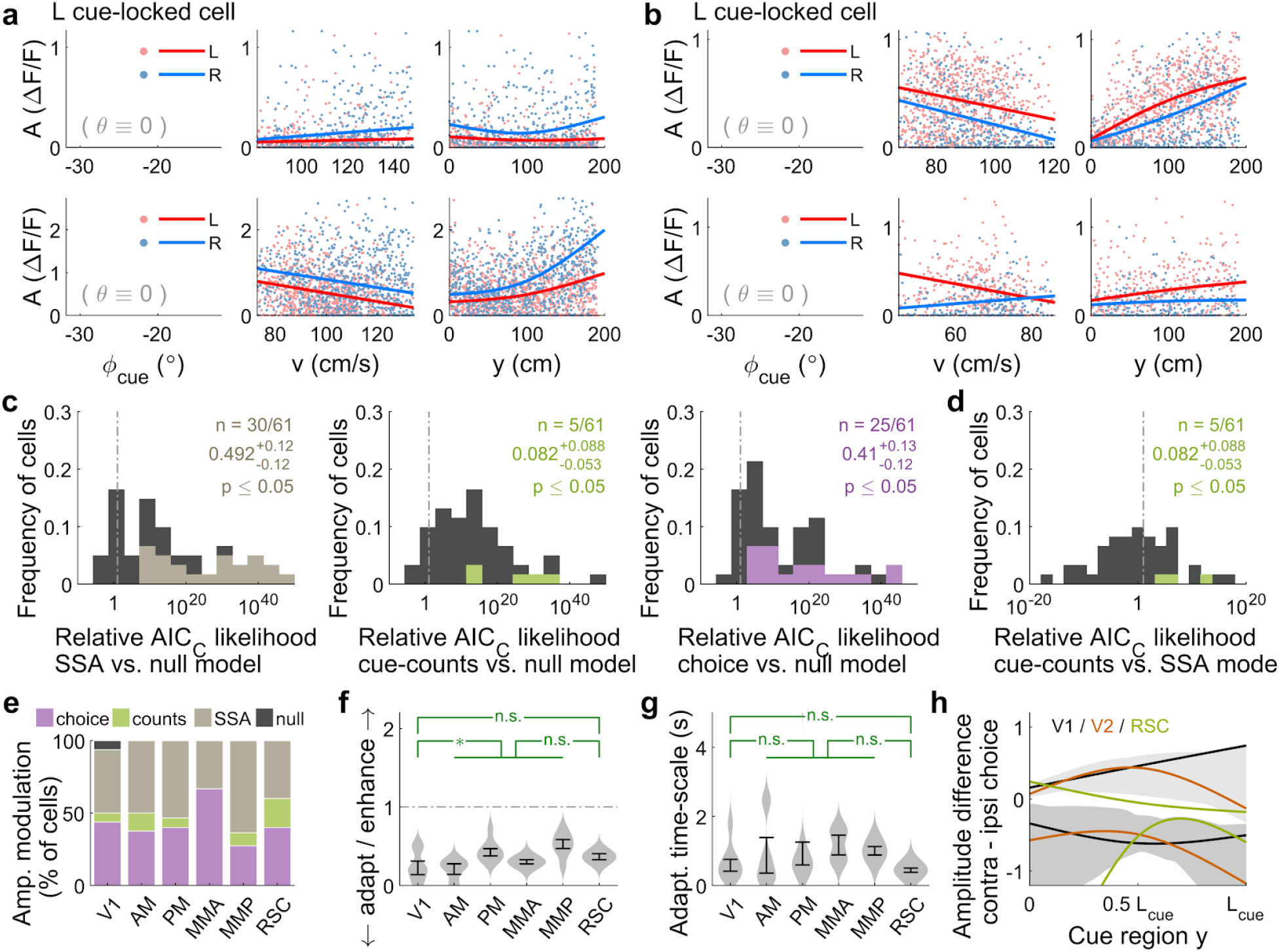
Qualitatively similar cue-locked amplitude modulations in control experiments with view angle restricted to be zero in the cue region. **(a-b,e-f,h)** As in Fig. 4, except using data from the control experiments. **(c-d,g)** As in Supplementary Fig. 5a-b,d, except using data from the θ-controlled experiments.

**Supplementary Figure 5.**
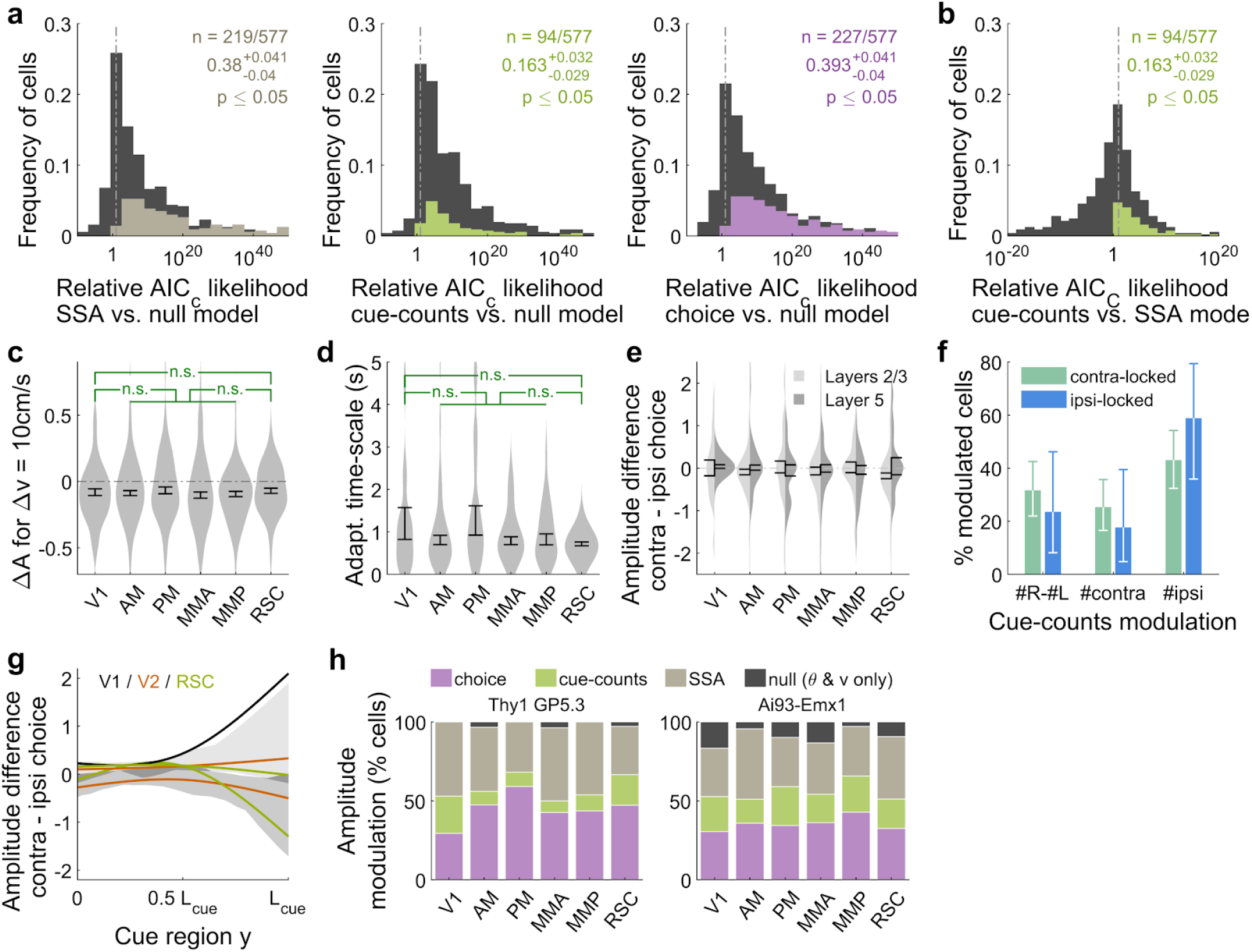
Additional statistics for amplitude modulations of cue-locked cells. **(a)** Distribution of AIC_C_ likelihood ratios for various amplitude-modulation models vs. the null hypothesis where cell responses only depend on an angular receptive field and speed. Colored areas corresponds to cells for which the indicated model is the best model for that cell (likelihood ratio n < 0.05). Data were pooled across all sessions. Note the logarithmic x-axis scale. **(b)** As in (a), distribution of AIC_C_ likelihood ratios for cue-counts model vs. the SSA model. Colored areas corresponds to cells for which the cue-counts model was the best model for that cell (likelihood ratio < 0.05). **(c)** Distribution (kernel density estimate) of predicted speed-induced changes in amplitudes for a change in speed of 10cm/s. Data were pooled across layers. Error bars: S.E.M. across cells. Stars: significant differences in means (Wilcoxon rank-sum test). **(d)** Distribution of adaptation/enhancement timescales for cells that favor the SSA model, defined as the time taken for the amplitude to recover to baseline by a factor of 1/*e*. Error bars: S.E.M. across cells. Stars: significant differences in means (Wilcoxon rank-sum test). **(e)** Distribution of choice modulation effect sizes for cue-locked cells in various areas/layers, defined as the maximum difference in predicted responses on contralateral- vs. ipsilateral-choice trials, divided by the mean response. Cells with numerically near-zero modulations (|δ*A^choice^*| ≤ 10^−4^) were excluded. Error bars: S.E.M. across cells. **(f)** Proportions out of all significantly cue-counts-modulated cells for which the best model is that which depends on the difference (left columns) in or single-side counts (middle and right columns) of cues, and shown separately for contralateral- and ipsilateral-cue-locked cells. Error bars: 95% C.I. across cells. **(g)** Choice modulation strength vs. location in the cue period for ipsilateral-cue-locked cells. **(h)** As in Fig. 4e, but for mice of two different strains. Data were pooled across layers.

**Supplementary Figure 6.**
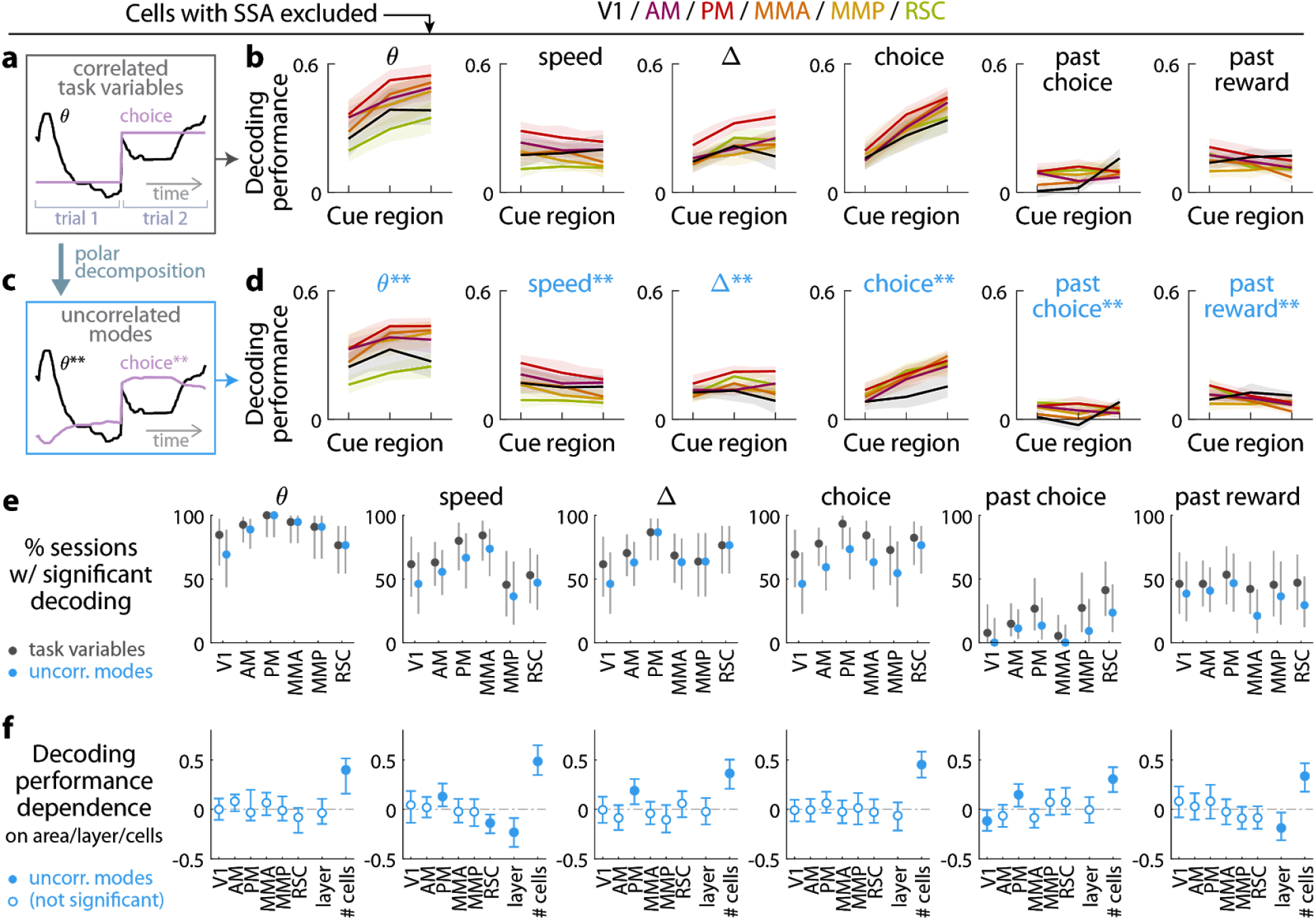
Evidence (and other task variables) can still be decoded from cue-locked response amplitudes, excluding cells that exhibit stimulus-specific adaptation (SSA). **(a-f)** As in Fig. 3, but with cells that favored the SSA model excluded from the data.

**Supplementary Figure 7.**
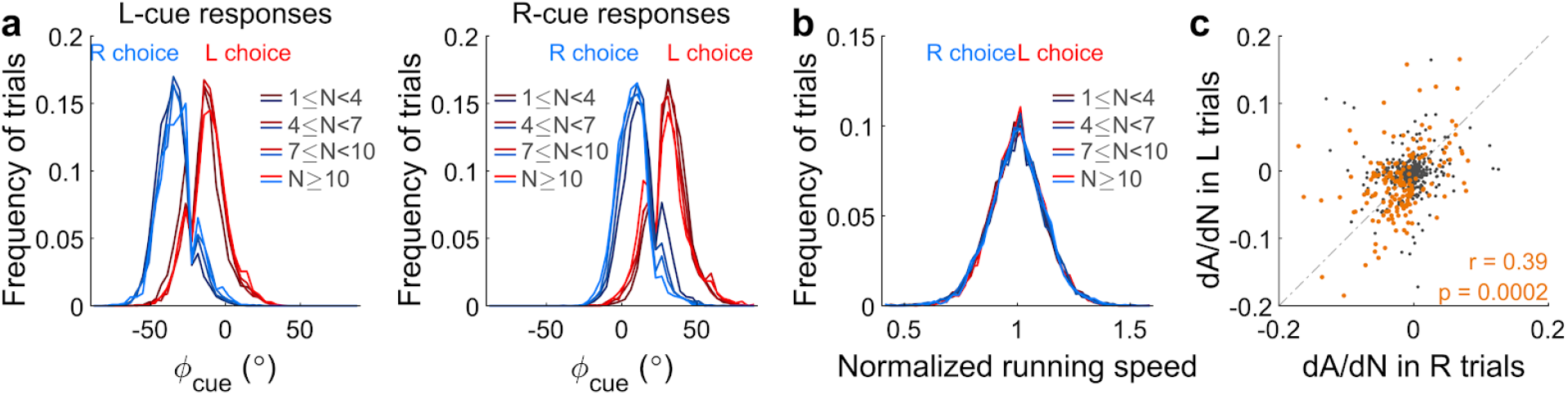
Statistics for evidence modulations of cue-response amplitudes, restricted to responses to the last cue in the last third of the cue region. **(a)** Distributions of visual angles at which the last cue appeared, for various numbers of left (left plot) and right (right plot) cue counts, and for trials of either choice category (color). **(b)** Distribution of running speeds (normalized to the median across trials in the session) at the instant at which the last cue appeared, for various numbers of cue counts and trials of either choice category. **(c)** Many cue-locked cells have last-cue response amplitudes that depend on the cue count. This plot shows the slope of linear regression of last-cue response amplitudes vs. cue counts, computed for either right-choice (x-axis) or left-choice (y-axis) trials only. Each point corresponds to one cell, and in orange are cells for which either of these slopes are significantly different from zero (permutation test; see Methods).

**Supplementary Figure 8.**
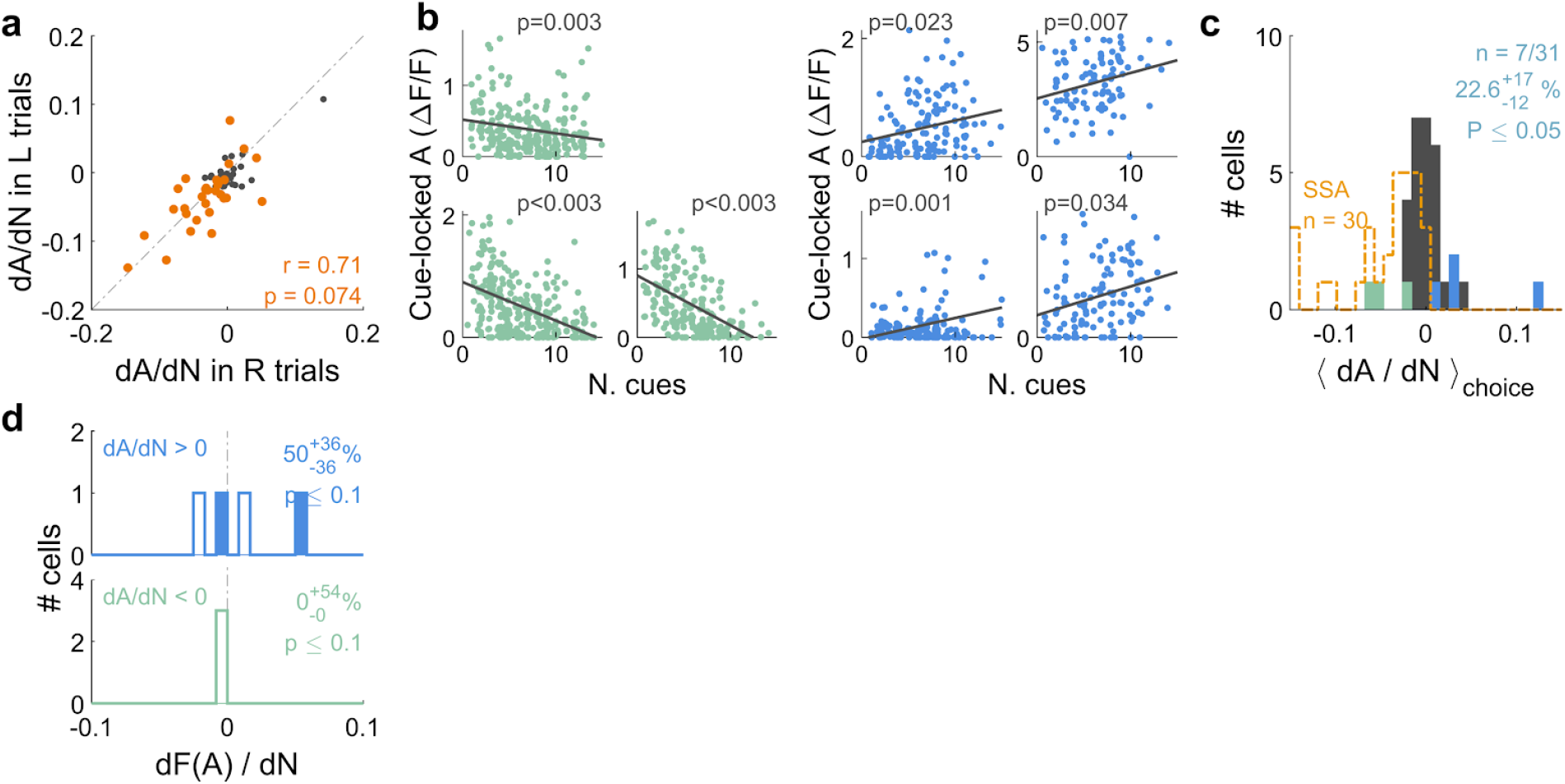
Qualitatively similar dependence of cue-locked response amplitudes on cue counts, in view-angle-locked control experiments. **(a)** As in Supplementary Fig. 7c, except for control experiments with view angle restricted to zero in the cue region. **(b-d)** As in Fig. 5c-f, except for control experiments with view angle restricted to zero in the cue region.

**Supplementary Figure 9.**
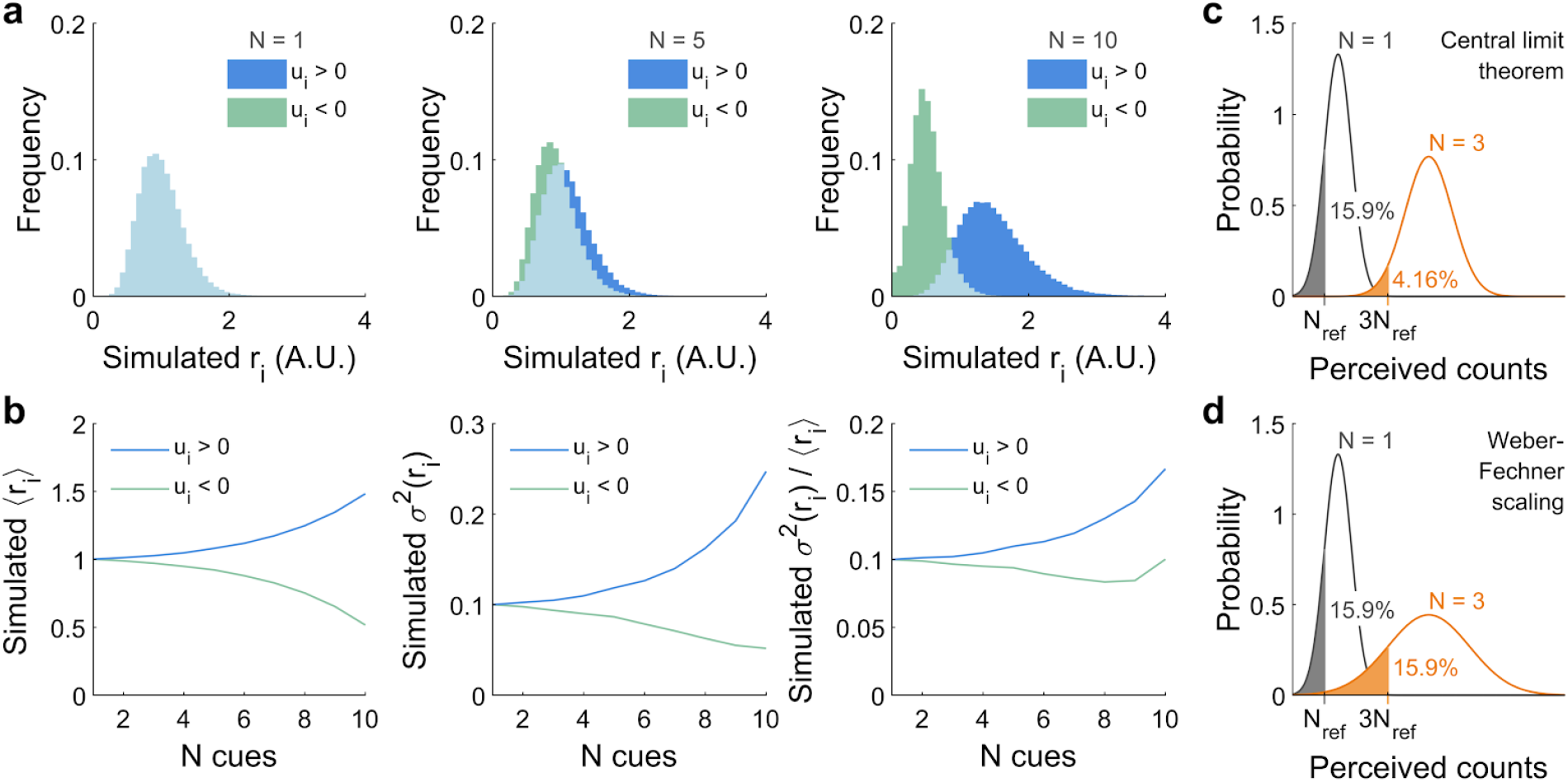
Illustration of sensory unit responses and count discrimination for different accumulator models. **(a)** Distribution of simulated sensory responses according to the feedback-loop model (*fdbk*), for a range of true counts (columns) that result in a stochastic distribution of accumulator states. Positively (negatively) modulated neurons are shown in blue (green). **(b)** Mean (left), variance (middle), and Fano factor (right plot) for the sensory responses in (a). **(c)** Illustration of how the distributions of perceived counts should change given a true count of *N* = 1 (gray) vs. a true count of *N* = 3 (orange), as predicted by the central limit theorem. Given 3 times more integrated counts, the width of the *N* = 3 distribution increases by 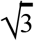 relative to that of the *N* = 1 distribution. This leads to less area falling under the reference level (15.9% for *N_ref_* as opposed to 4.16% for 3*N_ref_*), and corresponds to improved perceptual discriminability at higher counts. **(d)** As in (c), but as predicted by the Weber-Fechner Law of perceptual discrimination. This law prescribes that the *N* = 3 distribution is equivalent to taking the *N* = 1 distribution and scaling the perceived-counts axis by a factor of 3, including the reference level *N*_*ref*_ → 3*N*_*ref*_. This preserves the area under the reference level (15.9%) and for the same reason perceptual discriminability.

**Supplementary Figure 10.**
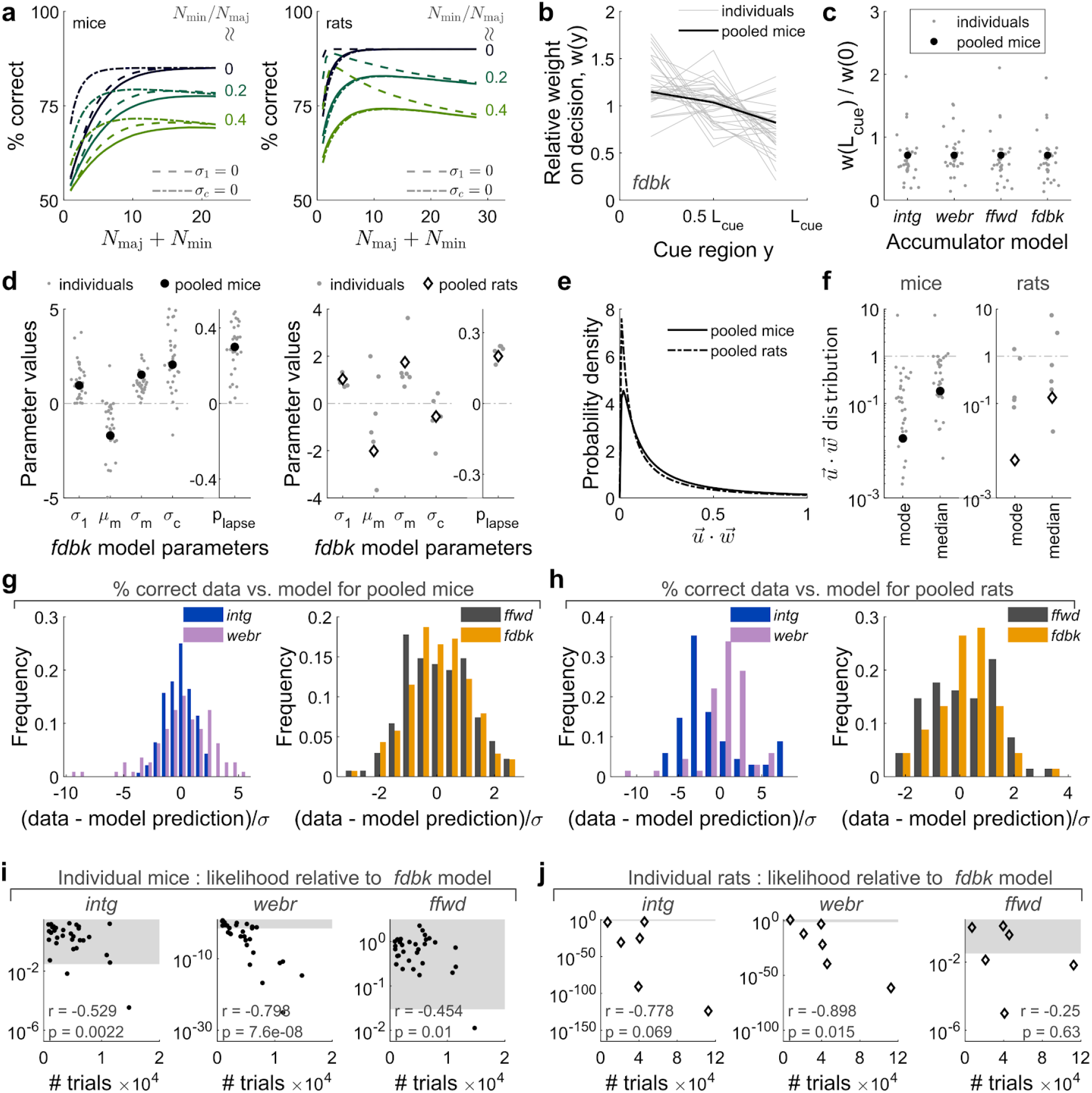
Additional goodness-of-fit metrics for psychophysical accumulator models. **(a)** Performance vs. counts as predicted by the *fdbk* model (solid lines), compared to the same when the sensory noise (dashed lines) or accumulator-comparison noise (dash-dotted lines) parameters were set to zero. The left plot is for fits to pooled mouse data, the right plot for fits to pooled rat data. **(b)** Spatial weights for evidence in the *fdbk* model for mice. These weights are highly comparable to and can be interpreted in the same way as Fig. 1f, with two differences. One, these models used much more behavioral data than the limited number of imaging sessions/mice in Fig. 1f. Two, theuse weights are in arbitrary units and therefore are expected to differ from those in Fig. 1f by an overall scaling of the y-axis in the two plots. **(c)** The same spatial weights in (b), but summarized by taking the ratio of weights at the end vs. start of the cue region. These weight ratios were highly similar across models (columns). **(d)** Optimized values for the free parameters of the *fdbk* model (except spatial weight parameters for mouse data, which are shown in (b-c)). **(e)** Modeled distribution of stochastic feedback-loop gains 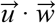 for the *fdbk* model fit to pooled mice and rat data. Each trial was one sample from this lognormal distribution, which had estimated parameters μ*m* and σ_*m*_ (see Methods) as shown in (d). **(f)** Mode and median for the feedback-loop gain distribution in (e), for various individual mice and rats (gray dots) as well as the pooled data (black disk for mice, open black diamond for rats). Only one data point fell outside of the range of this plot, being the mode statistic which had value 3.8 × 10−5 for one of the mice. **(g)** Distribution of residuals for accumulator model fits to pooled mouse data. Each entry in the histogram for a given model uses data with approximately the same *N*_*min*_/*N*_*maj*_ and *N*_*maj*_ + *N*_*min*_, and the residual is computed as the difference between the percent of correct trials in this data and the model prediction, divided by the estimated uncertainty (half of the 68% C.I.). A smaller residual (closer to 0) corresponds to a better fit. **(h)** As in (g), but for pooled rat data. **(i)** AIC_C_ likelihoods for various accumulator models relative to the fdbk model for individual mice, as in Fig. 6c, but as a function of the number of trials in the dataset for that mouse. *r*: Pearson’s correlation. Gray band: p-value threshold (same as Fig. 6c) below which model goodness-of-fit are not significantly different from that of the best model, at an α = 0.05 test level after correcting for multiple comparisons using the Benjamini-Hochberg procedure(Benjamini and Hochberg 1995). **(j)** As in (i), but for individual rats.

**Supplementary Figure 11.**
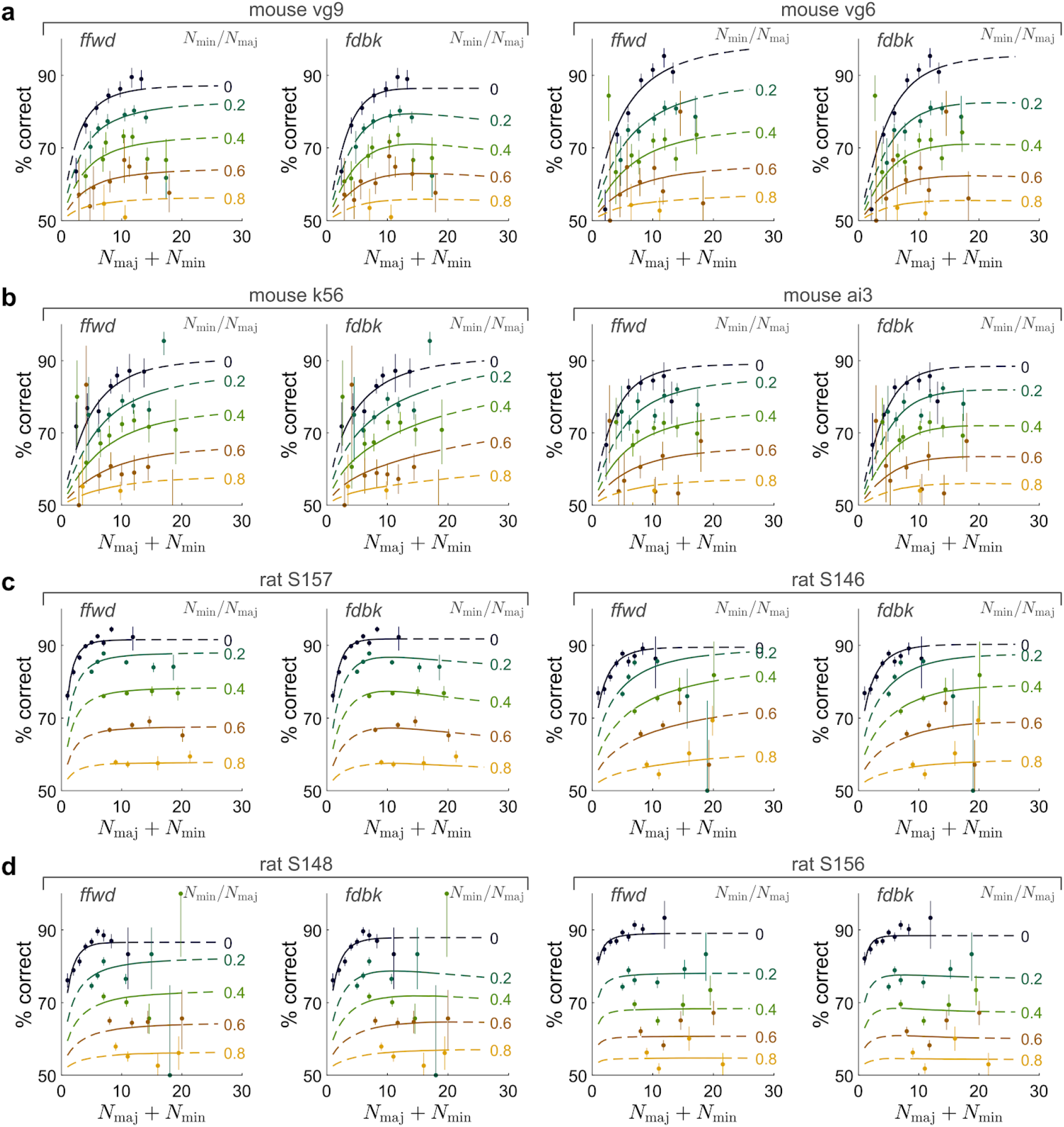
Data vs. accumulator model fits for individual mice and rats with the most number of behavioral trials. **(a-b)** as in Fig. 6e-f.

## Notes

### Competing Interest Statement

The authors have declared no competing interest.

